# Tumor Microenvironment Modulates Lineage Plasticity in Lung Squamous Cell Carcinoma

**DOI:** 10.64898/2026.06.22.733895

**Authors:** Yusuke Fujibayashi, Hiroyuki Ogawa, Quan Li, Roya Navab, Takamasa Koga, Yoshiaki Inoue, Nhu-An Pham, Hironori Hinokuma, Nicholas Bernards, Tadashi Sakane, Ko Matsumura, Yoshihisa Hiraishi, Fumi Yokote, Takahiro Yanagihara, Takashi Aoi, Yoshimasa Maniwa, Nikolina Radulovich, Ming-Sound Tsao, Kazuhiro Yasufuku

## Abstract

Lung squamous cell carcinoma (LUSC) is the second most common type of lung cancer, yet therapeutic options remain limited. A deeper understanding of its biology and molecular pathogenesis is essential for developing new treatment strategies. Here, we investigated the mechanisms of phenotypic plasticity in LUSC by comparing organoid-derived orthotopic lung models (ODOLs) and subcutaneous xenograft models (ODXs). ODXs showed greater tumor growth, squamous differentiation, and extracellular matrix (ECM) organization compared to ODOLs. Transcriptomic analyses revealed upregulation of multiple HIF1α and SOX2 target genes together with enhanced hypoxia signaling in ODXs. CRISPR/Cas9-mediated *HIF1α*-knockout ODXs showed reduced SOX2 expression, tumor growth, and ECM organization, whereas *SOX2*-knockout ODXs reduced tumor growth without affecting HIF1α and ECM organization. These results indicate that HIF1α regulates squamous lineage maintenance through SOX2 and ECM remodeling. Spatial transcriptomics revealed enrichment of basal cell-like and squamous-differentiated tumor states in ODXs, whereas ODOLs displayed less differentiated phenotypes. These findings identify the tumor microenvironment as a critical determinant of lineage plasticity in LUSC and provide mechanistic insight into how hypoxia shapes tumor differentiation.

## Introduction

Lung cancer is the common cause of cancer and cancer deaths worldwide ^1^. Non-small cell lung cancer (NSCLC) accounts for 80% of all lung cancers, with adenocarcinoma comprising 50% and squamous cell carcinoma accounting for 30% ^2^. Epidemiological studies indicated that lung squamous cell carcinoma (LUSC) has a poorer prognosis than lung adenocarcinoma (LUAD) ^3,4^. Nevertheless, efforts to investigate LUSC biology and therapeutic strategies have lagged those for LUAD.

Lung cancer organoids (LCOs) have recently emerged as a novel preclinical model in lung cancer research ^5^. LCOs retain the genetic and epigenetic alterations of the original patient tumors and are increasingly used to study cancer cell biology, tumorigenesis, and drug sensitivity and resistance mechanisms ^5–7^. Despite its potential, LCOs have some limitations. In human cancer tissue, cancer cells are surrounded by non-cancer cells such as fibroblasts, endothelial cells, immune cells, and extracellular matrix (ECM), which constitute the tumor microenvironment (TME). The TME has been demonstrated to play critical roles in cancer progression and treatment response ^8^. However, simple LCO models lack TME. To overcome this limitation, we have established a new orthotopic model by implanting LCOs into the lungs of mice and investigated the impact of TME on LUSC.

An orthotopic lung cancer model is more challenging to establish compared to the subcutaneous one because it requires substantially more technical expertise and resources for follow-up of the mice ^9^. However, orthotopic allows tumors to grow in an environment close to their native TME. In a subcutaneous model using human lung cancer cell lines, tumors often exhibit relatively uniform histology with reduced tissue polarity after inoculation ^10,11^. In contrast, LCOs maintain phenotypic characteristics that more closely resemble the original tumors when transplanted into mice ^12^. Although an orthotopic lung model established using conventional cell lines has been reported, the method using LCOs has been minimally investigated ^9,13,14^.

In this study, we have established the optimal method for generating organoid-derived orthotopic lung (ODOL) models for LUSC and compared ODOLs and subcutaneously grown organoid-derived xenograft (ODX) tumors. We showed that ODXs exhibited high SRY-box transcription factor 2 (SOX2) and a greater tumor growth ability, whereas ODOLs showed reduced SOX2 and lower growth rate. We also found that hypoxia-inducible factor 1-alpha (HIF1α), a major factor regulated by hypoxia in the TME, was highly expressed in ODXs. We further investigated how the TME influences LUSC lineage adaptations in LCOs *vs.* ODOLs or ODXs by spatial transcriptomics profiling, to gain new insights into the relationship between the TME and lineage plasticity in LUSC.

## Results

### Generation of ODOL model

Xenograft-derived organoid (XDO) model, XDO377, was established from LUSC-patient-derived xenograft (PDX) model PHLC377-X (**Figure 1A**). ODOLs were generated only by implantation of intact organoid structures. The establishment rates for ODOL-377 were correlated with the number of organoid cells implanted: 70.0% (7/10) with 2 million cells and 41.7% (5/12) with 1 million cells (**Figure 1B**). Implantation of dissociated single organoid cells at 2 million (7 mice) or 1 million (4 mice) failed to generate ODOL tumors. Furthermore, 2 million cells as intact organoids engrafted significantly larger tumors than with other conditions (p<0.0001) (**Figure 1B-D**).

**Figure 1.**
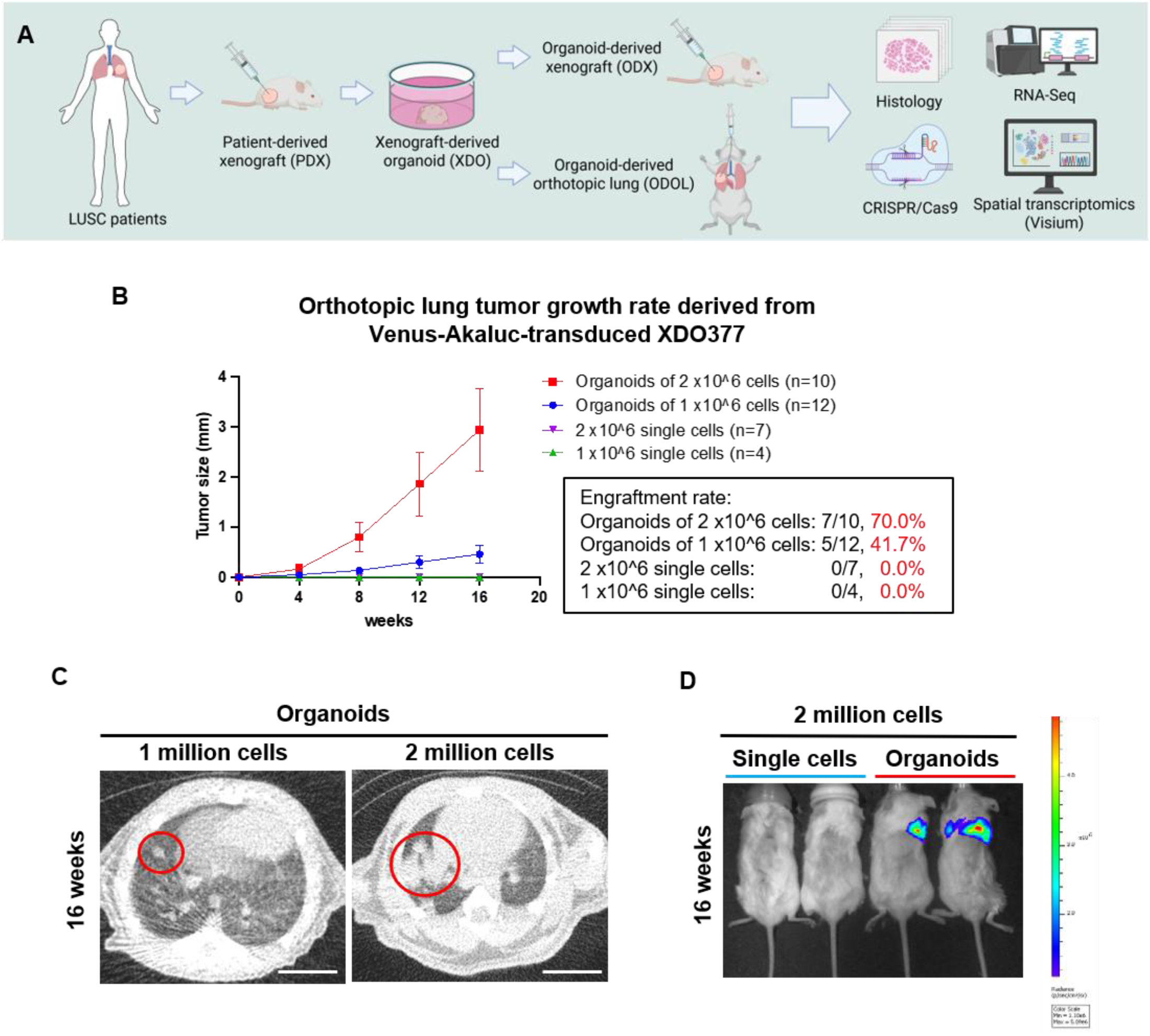
Establishment of an ODOL-377. (A) The overall study schema of the project. (B) The growth rate of orthotopic lung tumors derived from Venus-Akaluc-transduced XDO377, which are presented by dividing the groups into 1 x 10^6^ or 2 x 10^6^ cells, either as intact organoids or single-cell. The engraftment rate for each group is shown. (C) CT images of ODOL-377 of 1 X10^6^ cells or 2 x10^6^ cells with preserved organoid structures. Two million cells of organoids grew larger tumor than 1 X10^6^ cells in 16 weeks. Scale bar is 2 mm. (D) The Akaluc signal was detected in the ODOL-377 with organoids of 2 x10^6^ cells, but not in the model transplanted with single-cell.

### Tumor differentiation and proliferation in ODOL and ODX models

Another XDO model, XDO149, was established from LUSC-PDX model PHLC149-X. To facilitate tumor identification, red fluorescence protein (RFP) was transduced into XDO377 (XDO377^RFP^) and XDO149 (XDO149^RFP^) (**Figure 2A**). ODOLs using XDO377^RFP^ (ODOL-377^RFP^) were successfully established in 15/34 cases (44.1%), and ODXs using XDO377^RFP^ (ODX-377^RFP^) were successfully established in 4/4 cases (100%) **(Figure 2B, 2C)**. ODOLs using XDO149^RFP^ (ODOL-149^RFP^) were successfully established in 3/12 cases (25.0%), and ODXs using XDO149^RFP^ (ODX-149^RFP^) were successful in 5/5 cases (100%). Both ODX-377^RFP^ and ODX-149^RFP^ formed well-differentiated keratinizing LUSC, while ODOL-377^RFP^ and ODOL-149^RFP^ showed poorer differentiation (**Figure 2D**). By immunohistochemistry (IHC), ODX-377^RFP^ and ODX-149^RFP^ were positive for SOX2 and p40, consistent with the staining patterns of the parental patient-derived xenografts (PDXs) and XDOs (**Figure 2E**) (**Supplementary Figure 1)**. In contrast, ODOL-377^RFP^ and ODOL-149^RFP^ exhibited significantly lower expression of SOX2 and p40 compared to ODX-377^RFP^ and ODX-149^RFP^ **(Figure 2E, 2F)**. ODX-377^RFP^ and ODX-149^RFP^ showed increased collagen organization by Picrosirius red staining compared with ODOL-377^RFP^ and ODOL-149^RFP^ **(Supplementary Figure 2)**. Moreover, ODOL-377^RFP^ and ODOL-149^RFP^ showed significantly lower growth rate compared to ODX-377^RFP^ and ODX-149^RFP^ (**Figure 2G**). These results suggest that different TMEs between the subcutaneous and lung tissue may influence the differentiation of LUSC.

**Figure 2.**
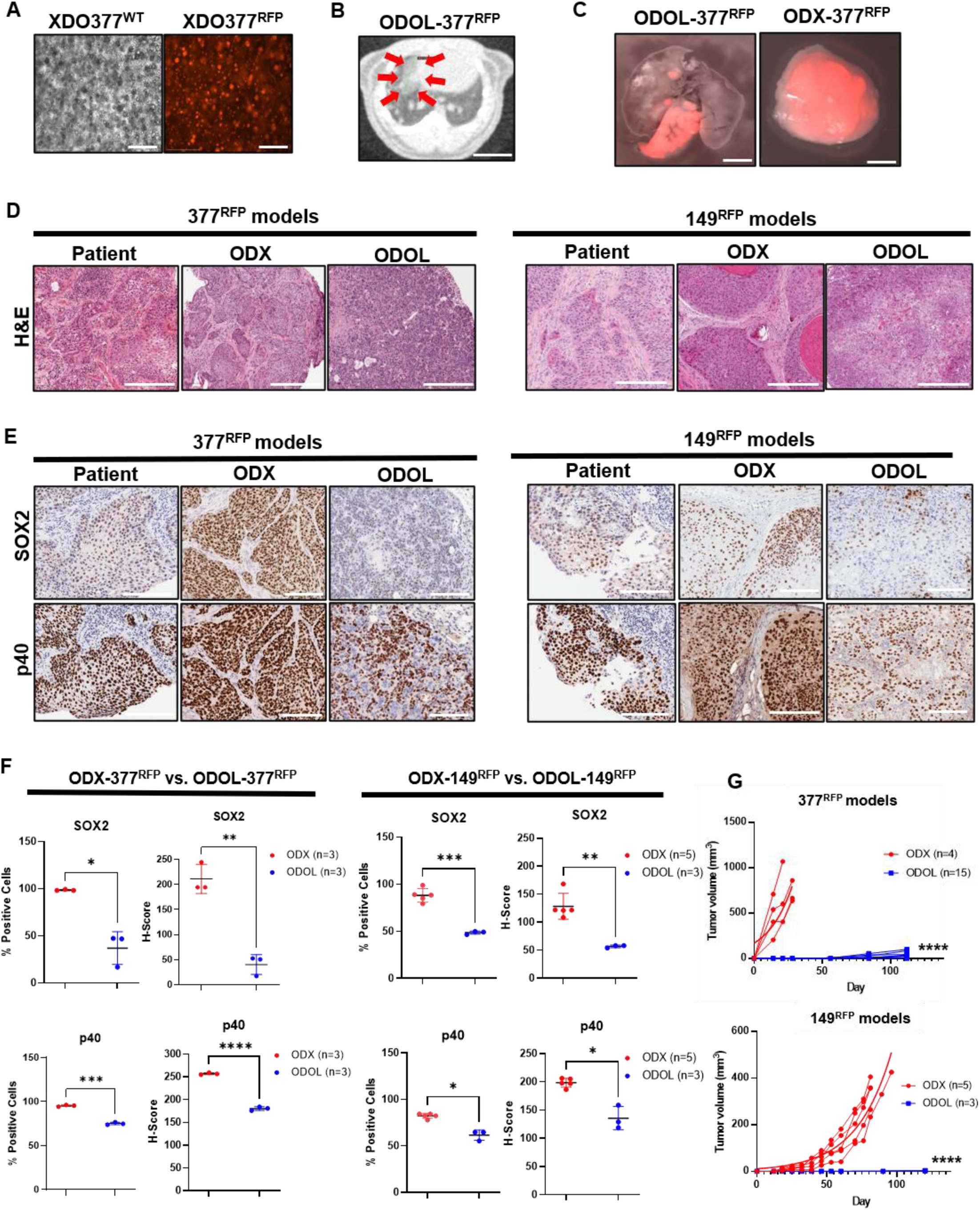
Tumor differentiation and proliferation in ODOL and ODX models. (A) Confocal micrographs of XDO377 before and after transduction with RFP. (B) CT image of ODOL-377^RFP^. Red arrows indicate the location of tumor. (C) The RFP in ODOL (left image) and ODX (right image) were confirmed using macroscopy. (D) H&E staining in 377^RFP^ and 149^RFP^ models (Patient tumors, ODXs, and ODOLs). (E) IHC staining for p40 and SOX2 in 377^RFP^ and 149^RFP^ models (Patient tumors, ODXs, and ODOLs). (F) The staining intensity in Figure 1E was quantitatively evaluated using % positive stained cells and H-score with HALO software. (G) Tumor growth rate is shown between ODXs and ODOLs. Scale bars are 65 μm (A), 1mm (B), 2mm (C), 300 μm (D), 200 μm (E).

### TME modulates phenotypic changes in LUSC

To further investigate the impact of orthotopic versus subcutaneous TME, we performed whole exome/shallow genome sequencing (WES/sWGS) and RNA sequencing (RNA-seq) on ODX-377^RFP^ and ODOL-377^RFP^.

WES/sWGS demonstrated high concordance of both mutations and gene copy number among XDO377^RFP^, ODX-377^RFP^, and ODOL-377^RFP^ **(Figure 3A, 3B)**. Clonal evolution analysis also showed same "branched" evolution pattern and similar clone prevalence across the models (**Figure 3C**). By RNA-seq, differentially expressed genes (DEGs) analysis showed that 618 genes were upregulated and 1,788 genes downregulated in ODX-377^RFP^.

**Figure 3.**
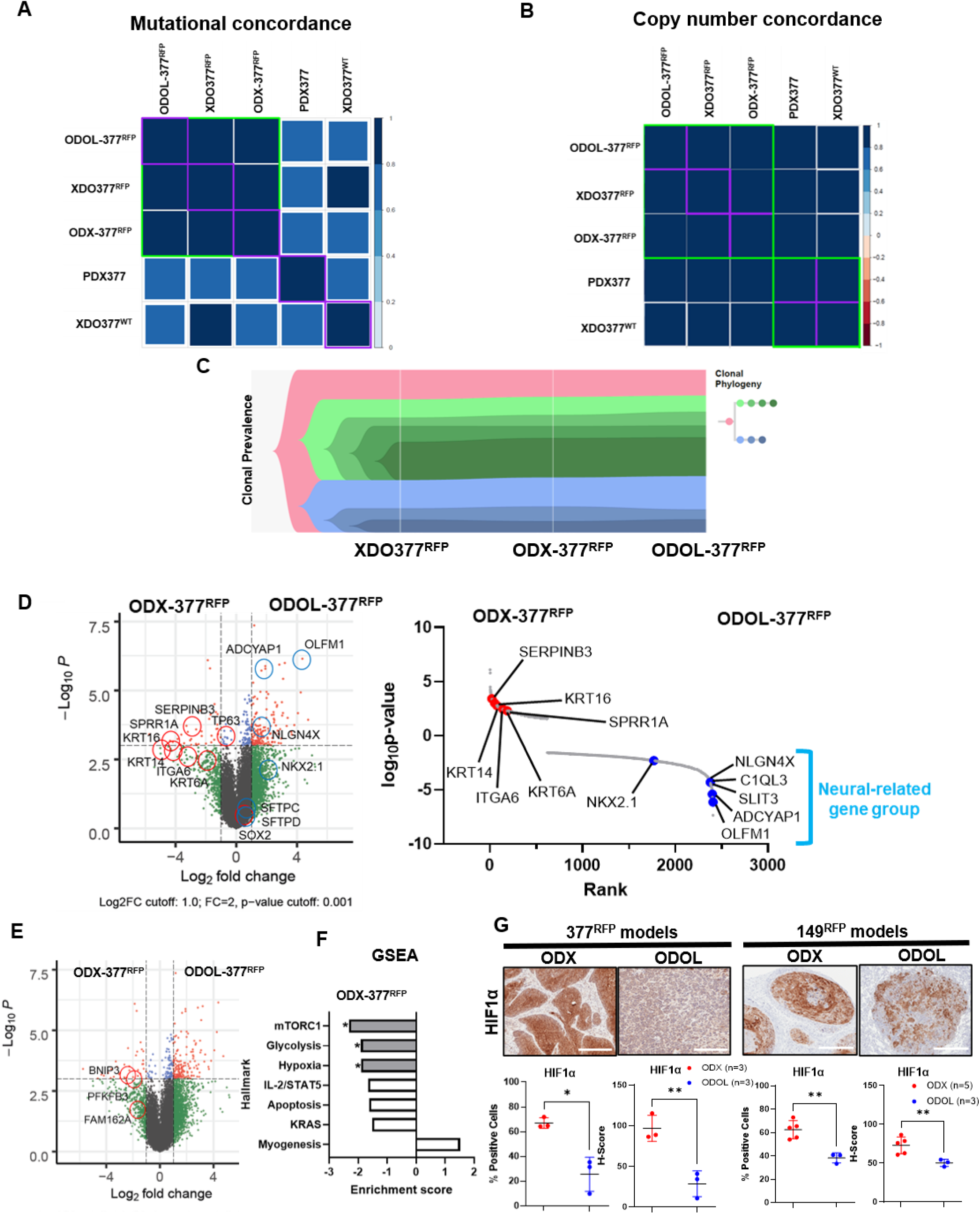
TME influences phenotypic changes in LUSC. (A) Mutational concordance heatmap among XDO377^RFP^, ODX-377^RFP^, and ODOL-377^RFP^. (B) Copy number concordance heatmap among XDO377^RFP^, ODX-377^RFP^, and ODOL-377^RFP^. (C) Clonal prevalence among XDO377^RFP^, ODX-377^RFP^, and ODOL-377^RFP^. (D) The left figure is the volcano plot of DEGs between ODX-377^RFP^ and ODOL-377^RFP^. The right figure is the Rank-log10 p-value plot between ODX-377^RFP^ and ODOL-377^RFP^. (E) Volcano plot of DEGs between ODX-377^RFP^ and ODOL-377^RFP^. (F) GSEA between ODX-377^RFP^ and ODOL-377^RFP^. (G) IHC staining for HIF1α between ODX-377^RFP^/-149^RFP^ and ODOL-377^RFP^/-149^RFP^. Scale bars are 300 μm. The staining intensity was quantitatively evaluated using % positive stained cells and H-score with HALO software.

Among these RNAs, LUSC markers downstream of SOX2 such as Keratin (KRT) 6A, KRT14, KRT16, Small Proline-Rich Protein 1A (SPRR1A), and Serpin Family B Member 3 (SERPINB3) were upregulated in ODX-377^RFP^. In contrast, LUAD markers such as NK2 Homeobox 1 (NKX2.1) and neural-regulated genes, including Neuroligin 4 X-Linked (NLGN4X), Complement C1q Like 3 (C1QL3), Adenylate Cyclase Activating Polypeptide 1 (ADCYAP1), and Olfactomedin 1 (OLFM1), were upregulated in ODOL-377^RFP^ (**Figure 3D**). Furthermore, downstream genes of HIF1α, including BCL2 Interacting Protein 3 (BNIP3), 6-Phosphofructo-2-kinase/fructose-2,6-bisphosphatase 3 (PFKFB3), and Family with Sequence Similarity 162 Member A (FAM162A), were upregulated in ODX-377^RFP^ (**Figure 3E**). Gene set enrichment analysis (GSEA) using hallmark showed that mechanistic target of rapamycin (mTOR), glycolysis, and hypoxia were upregulated in ODX-377^RFP^ (**Figure 3F**).

The HIF1α protein expression was significantly upregulated in both ODX-377^RFP^ and ODX-149^RFP^ compared to their respective ODOL tumors (**Figure 3G**). These findings suggest that HIF1α, a key condition of the TME influences phenotypic changes in LUSC, putatively via SOX2.

### HIF1α contributes to the maintenance of LUSC lineage

When XDO377^RFP^ and XDO149^RFP^ were cultured under a hypoxic condition, HIF1α, SOX2, and Delta Np63 protein levels were significantly elevated compared to in a normoxic condition. SOX2 and HIF1α were also significantly upregulated in ODX-377^RFP^ compared to ODOL-377^RFP^ (**Figure 4A**) (**Supplementary Figure 3A)**. The mRNA expression was consistent with the changes at protein level in XDOs **(Supplementary Figure 3B)**.

**Figure 4.**
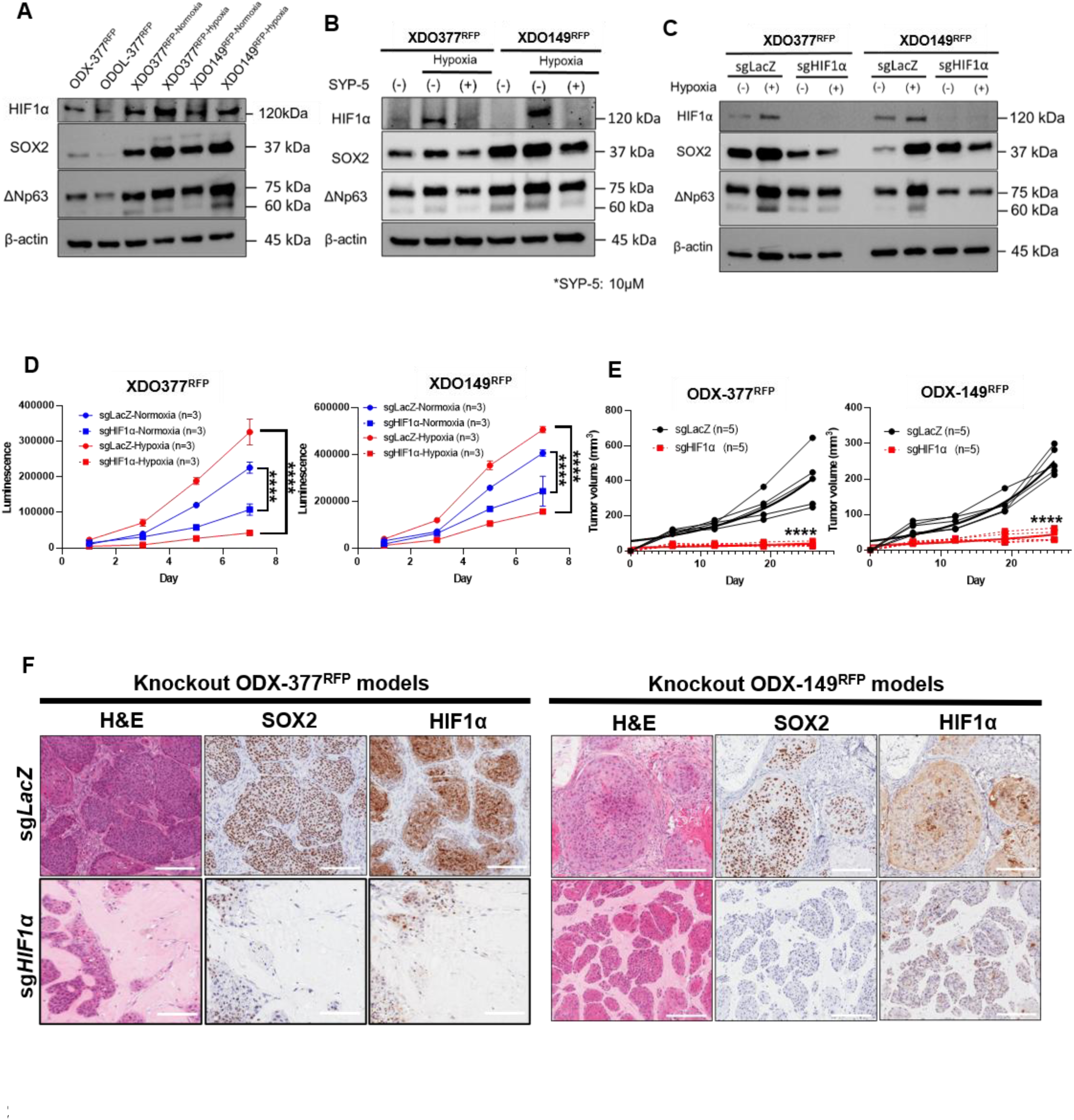
HIF1α contributes to the maintenance of LUSC lineage. (A) Western blotting: ODX-377^RFP^ vs ODOL-377^RFP^; normoxic XDOs vs hypoxic XDOs. (B) Western blotting: XDO377^RFP^ and XDO149^RFP^ under SYP-5. (C) Western blotting: XDO377^RFP-sg*LacZ*^/XDO^149RFP-sg*LacZ*^ vs XDO377^RFP-sg*HIF1α*^/XDO149^RFP-sg*HIF1α*^ under a normoxic and hypoxic condition. (D) The cell proliferation: XDO377^RFP-sg*LacZ*^/XDO149^RFP-sg*LacZ*^ vs XDO377^RFP-sg*HIF1α*^/XDO149^RFP-sg*HIF1α*^ under normoxic and hypoxic condition. (E) Tumor growth rates: ODX-377^RFP-sg*LacZ*^/-149^RFP-sg*LacZ*^ vs ODX-377^RFP-sg*HIF1α*^/-149^RFP-sg*HIF1α*^. (F) H&E and IHC staining for SOX2 and HIF1α between ODX-377^RFP-sg*LacZ*^/-149^RFP-sg*LacZ*^ vs ODX-377^RFP-sg*HIF1α*^/-149^RFP-sg*HIF1α*^. Scale bars are 200 μm.

An HIF1 inhibitor SYP-5 (SML1894, Sigma) significantly suppressed SOX2 and Delta N p63 expression under hypoxia (**Figure 4B**) (**Supplementary Figure 3C)**. When HIF1α was knocked-out by CRISPR/Cas9 and sg*HIF1α* in the XDOs, SOX2 and Delta Np63 expression were suppressed under hypoxia (**Figure 4C**). The XDO377^RFP-sg*HIF1α*^ and XDO149^RFP-sg*HIF1α*^ also showed significantly lower proliferation (**Figure 4D**), and in the mouse subcutaneous tissue, the ODX-377^RFP-sg*HIF1α*^ and ODX-149^RFP-sg*HIF1α*^ demonstrated significantly growth retardation (**Figure 4E**) (**Supplementary Figure 4)**. These results suggest that HIF1α is a strong regulator of proliferation in LUSC ODX cells, both *in vitro* and *in vivo*.

By IHC, ODX-377^RFP-sg*HIF1α*^ and ODX-149^RFP-sg*HIF1α*^ showed lower HIF1α and SOX2 levels (**Figure 4F**, **Supplementary Figure 5A)**. However, p40 expression was not significantly altered **(Supplementary Figure 5B)**. RNA-seq analyses showed keratinization was significantly downregulated in ODX-377^RFP-sg*HIF1α*^ and ODX-149^RFP-sg*HIF1α*^ (**Figure 5A**, **5C)**. Consistent with this, hematoxylin and eosin (H&E) staining of ODX-377^RFP-sg*HIF1α*^ and ODX-149^RFP-sg*HIF1α*^ showed a lack of keratinization. Immunofluorescence (IF) staining confirmed that the high expression of SOX2 and HIF1α was present in the same cells **(Supplementary Figure 6)**. Overall, our results indicate that HIF1α, which was activated by hypoxia in subcutaneous environment, upregulates SOX2 expression and keratinization, thereby regulating the differentiation and tumor growth in LUSC.

**Figure 5.**
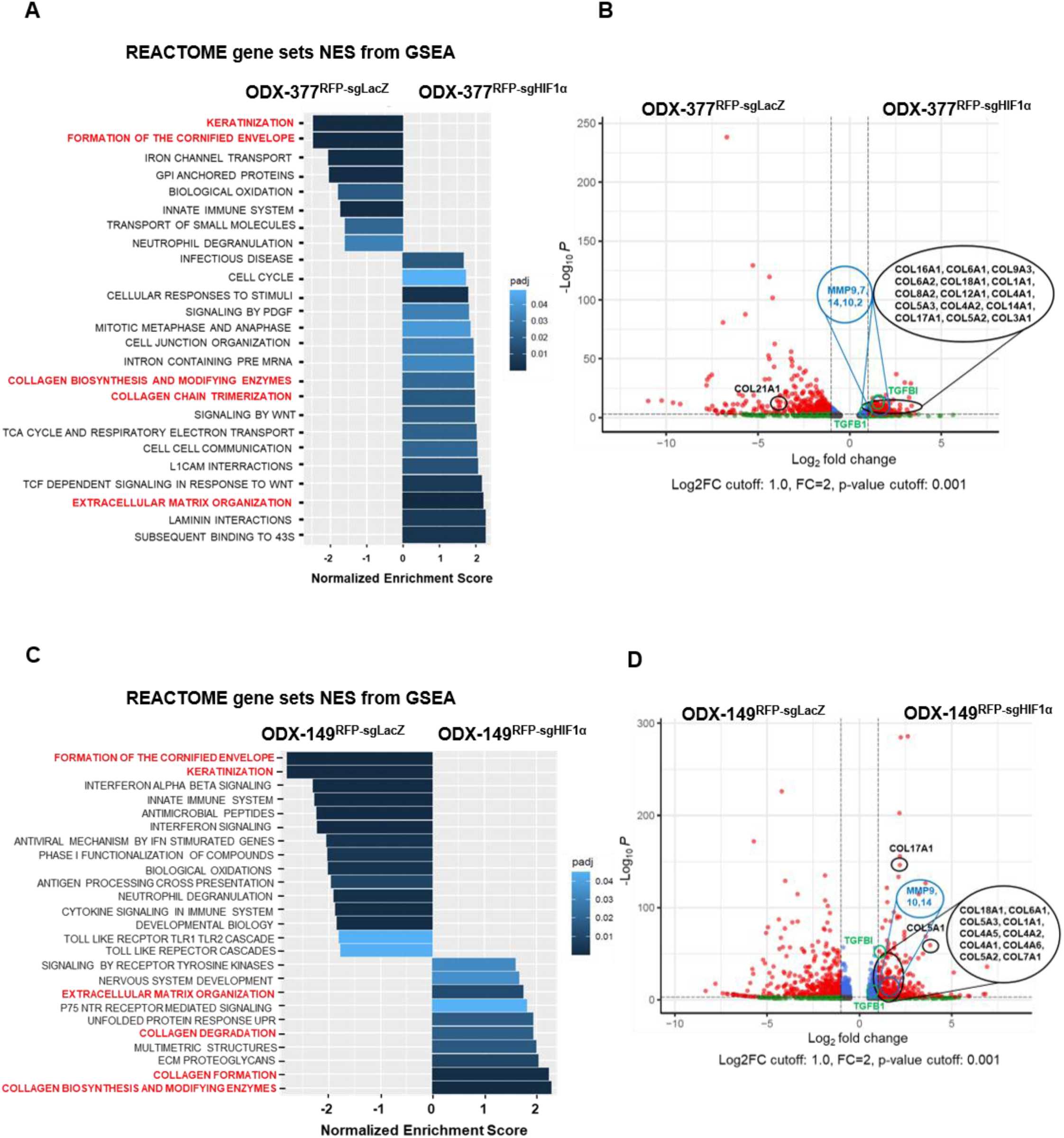
HIF1α regulates keratinization, epithelial-mesenchymal transition (EMT), and extracellular matrix remodeling in LUSC. (A) GSEA using REACTOME between ODX-377^RFP-sg*LacZ*^ vs ODX-377^RFP-sg*HIF1α*^. (B) Volcano plot of DEGs between ODX-377^RFP-sg*LacZ*^ vs ODX-377^RFP-sg*HIF1α*^. (C) GSEA using REACTOME between ODX-149^RFP-sg*LacZ*^ vs ODX-149^RFP-sg*HIF1α*^. (D) Volcano plot of DEGs between ODX-149^RFP-sg*LacZ*^ vs ODX-149^RFP-sg*HIF1α*^.

### HIF1α regulates epithelial-mesenchymal transition (EMT) and extracellular matrix remodeling in LUSC

Principal component analysis (PCA) and concordance rate analyses of ODX-377^RFP-sg*LacZ*^ and ODX-149^RFP-sg*LacZ*^ showed markedly different genetic profiles between the two ODX models, reflecting heterogeneity in LUSC **(Supplementary Figure 7A, 7B)**. GSEA analysis of the RNA-seq data using hallmark also showed significant upregulation of epithelial-mesenchymal transition (EMT) in ODX-377^RFP-sg*HIF1α*^ and ODX-149^RFP-sg*HIF1α*^ **(Supplementary Figure 7C, 7D)**. DEGs analysis revealed many collagen genes were upregulated in ODX-377^RFP-sg*HIF1α*^ and ODX-149^RFP-sg*HIF1α*^ (**Figure 5B**, **5D)**. Furthermore, transforming growth factor beta induced protein (TGFI) and many matrix metalloproteinases (MMPs) were upregulated in ODX-377^RFP-sg*HIF1α*^ and ODX-149^RFP-sg*HIF1α*^ (**Figure 5B**, **5D)**. However, picrosirius red staining revealed non-organized collagen in ODX-377^RFP-sg*HIF1α*^ and ODX-149^RFP-sg*HIF1α*^, which is confirmed by second harmonic generation (SHG) analysis **(Supplementary Figure 8)**. Therefore, the results suggest that despite a robust collagen synthesis, elevated MMPs activity may lead to the degradation of collagen fibers, thereby hindering organized matrix assembly. These findings suggest that HIF1α may also regulate LUSC lineage by remodeling ECM, particularly collagen organization. Collectively, these results support the hypothesis that the TME modulates lineage plasticity in LUSC.

### The role of SOX2 in the LUSC lineage

Two *SOX2*-knockout (sg*SOX2*) XDOs, XDO377^RFP-sg*SOX2*^ and XDO149^RFP-sg*SOX2*^, were established. A significantly lower proliferation was found in XDO377^RFP-sg*SOX2*^ and XDO149^RFP-sg*SOX2*^ (**Figure 6A**). While SOX2 protein expression was entirely inapparent in XDO377^RFP-sg*SOX2*^, minimal residual SOX2 expression was detected and with a band at a lower molecular weight in XDO149^RFP-sg*SOX2*^ (**Figure 6B**). When XDO377^RFP-sg*SOX2*^ and XDO149^RFP-sg*SOX2*^ were cultured under hypoxia, SOX2 expression remained completely suppressed in XDO377^RFP-sg*SOX2*^, but not in XDO149^RFP-sg*SOX2*^, which showed an increase expression of the lower molecular weight band (**Figure 6B**). Taking into account the proliferation and the nucleotide sequence analysis of SOX2 gene in XDO149^RFP-sg*SOX2*^ **(Supplementary Figure 9A)**, this lower molecular weight SOX2 was most likely nonfunctional. In XDO377^RFP-sg*SOX2*^ and XDO149^RFP-sg*SOX2*^, HIF1α expression was elevated under hypoxia. This result indicates that HIF1α acts upstream of SOX2.

**Figure 6.**
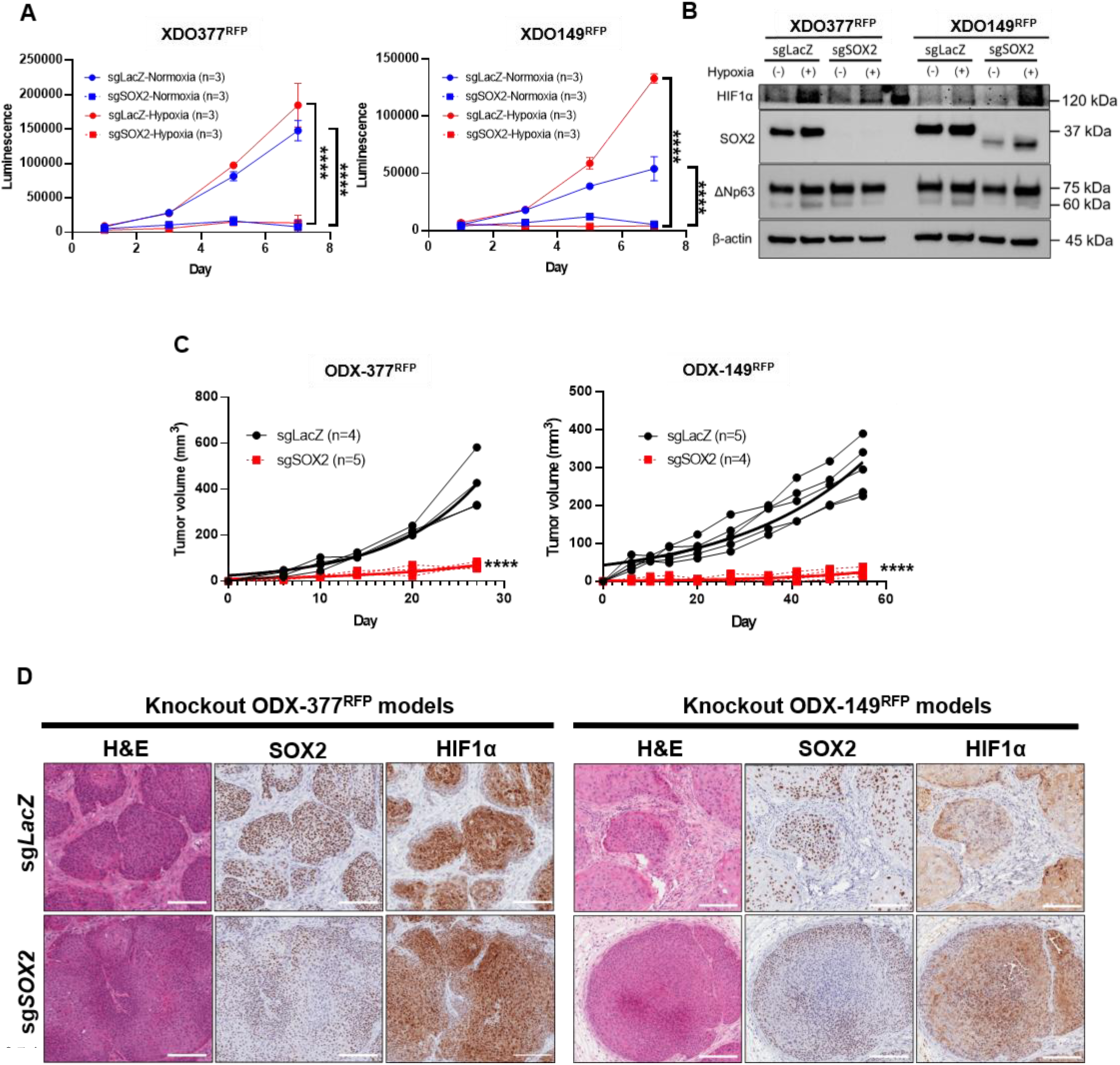
Role of SOX2 in the LUSC lineage. (A) The cell proliferation: XDO377^RFP-sg*LacZ*^/XDO149^RFP-sg*LacZ*^ vs XDO377^RFP-sg*SOX2*^/XDO149^RFP-sg*SOX*2^. (B) Western blotting: XDO377^RFP-sg*LacZ*^/XDO149^RFP-sg*LacZ*^ vs XDO377^RFP-sg*SOX2*^/XDO149^RFP-sg*SOX2*^ under a normoxic and hypoxic condition. (C) Tumor growth rates are shown between ODX-377^RFP-sg*LacZ*^/-149^RFP-sg*LacZ*^ vs ODX-377^RFP-sg*SOX2*^/-149^RFP-sg*SOX2*^. (D) H&E and IHC staining for SOX2 and HIF1α between ODX-377^RFP-sg*LacZ*^/-149^RFP-sg*LacZ*^ vs ODX-377^RFP-sg*SOX2*^/-149^RFP-sg*SOX2*^. Scale bars are 200 μm.

In subcutaneous tissue of mice, both ODX-377^RFP-sg*SOX2*^ and ODX-149^RFP-sg*SOX2*^ showed significantly lower growth rate (**Figure 6C**) (**Supplementary Figure 4)**. By IHC, ODX-377^RFP-sg*SOX2*^ and ODX-149^RFP-sg*SOX2*^ showed significantly reduced SOX2 staining but no significant change in HIF1α expression (**Figure 6D**) (**Supplementary Figure 10A)**. These results are consistent with the *in vitro* data (**Figure 6B**) and HIF1α acting upstream of SOX2. The expression of p40 was slightly decreased in ODX-377^RFP-sg*SOX2*^, whereas it was slightly increased in ODX-149^RFP-sg*SOX2*^, yielding inconsistent results **(Supplementary Figure 10B)**. With H&E and picrosirius red stains, ODX-377^RFP-sg*SOX2*^ and ODX-149^RFP-sg*SOX2*^ did not exhibit ECM remodeling as observed in ODX-377^RFP-sg*HIF1α*^ and ODX-149^RFP-sg*HIF1α*^ (**Figure 6D**) (**Supplementary Figure 8)**. These results reaffirm that SOX2 is an essential gene for LUSC cell survival and SOX2 is downstream of HIF1α, and HIF1α regulates ECM remodeling independent of SOX2.

### Re-establishment of XDOs reveals TME-dependent plasticity

We re-established organoids from ODX-377^RFP^ and ODOL-377^RFP^, respectively to assess the phenotypic plasticity. By RNA-seq, DEGs analysis showed that 2,406 genes were different between ODX-377^RFP^ and ODOL-377^RFP^ tumors, but only 44 genes were different between re-established organoids derived from ODX-377^RFP^ and ODOL-377^RFP^ **(Supplementary Figure 11A)**. PCA showed ODX-377^RFP^ and ODOL-377^RFP^ exhibited distinct directions of gene expression, however, the expressed genes of re-established organoids showed similar patterns to the original XDO377^RFP^ **(Supplementary Figure 11B)**. Taken together, the phenotypic changes of organoids are regulated by the TME, and removal of TME influences allows these changes to revert to their original state.

### Spatial transcriptomics revealed lineage shifts driven by cell type states

Spatial transcriptomics using Visium HD was performed for XDO377^RFP^/-149^RFP^, ODX-377^RFP^/-149^RFP^, and ODOL-377^RFP^/-149^RFP^. Both XDO377^RFP^ and XDO149^RFP^ were composed entirely of epithelial cells. In ODX-377^RFP^ and ODOL-377^RFP^, tumors were predominantly composed of epithelial cells, whereas tumors were largely composed of keratinized squamous epithelial cells in ODX-149^RFP^ and ODOL149^RFP^ **(Figure 7A, 7C) (Supplementary Figure 12A, 12D)**. This indicates significant heterogeneity exists in the lineage differentiation potential of LUSC organoid models, in term of keratinization, and this heterogeneity between LUSC models 377 and 149 is consistent with the RNA-seq results from sg*LacZ*-ODXs **(Supplementary Figure 7A, 7B)**. Furthermore, most tumor cells in ODX-377^RFP^ and ODX-149^RFP^ were basal cell-like, whereas the proportion of basal cell-like tumor cells decreased in ODOLs. In addition, alveolar type 1 (AT1)- and AT2-like tumor cells increased in ODOLs **(Figure 7A, 7C) (Supplementary Figure 12A, 12D)**. Pseudotime analysis showed that XDO followed a common trajectory, whereas ODX and ODOL exhibited distinct differentiation patterns. Furthermore, basal cell-like tumor cells were enriched along the trajectory toward ODX, while AT1- and AT2-like tumor cells were enriched along the trajectory toward ODOL **(Figure 7B, 7E) (Supplementary Figure 12B, 12E)**. These results suggest that the TME regulates tumor phenotypes by modulating tumor cell state plasticity. Furthermore, well-differentiated LUSC observed in ODXs and ODOLs is also supported by the higher expression of genes such as KRT5, KRT6A, KRT13, KRT14, KRT15, KRT16, KRT17, SERPINB3, and SERPINB5 compared with organoids **(Supplementary Figure 12C, 12F)**. However, ODOLs exhibited lower expression of these squamous differentiation markers than ODXs. Downstream genes of HIF1α, FAM162A and PFKFB3, were also reduced in ODOLs. These results are consistent with those previously discussed. Overall, spatial transcriptomics confirmed the differences among organoids, ODXs, and ODOLs, and further revealed that the TME has a substantial impact on the LUSC lineage by regulating the plasticity between basal cells and AT1/AT2 cells.

**Figure 7.**
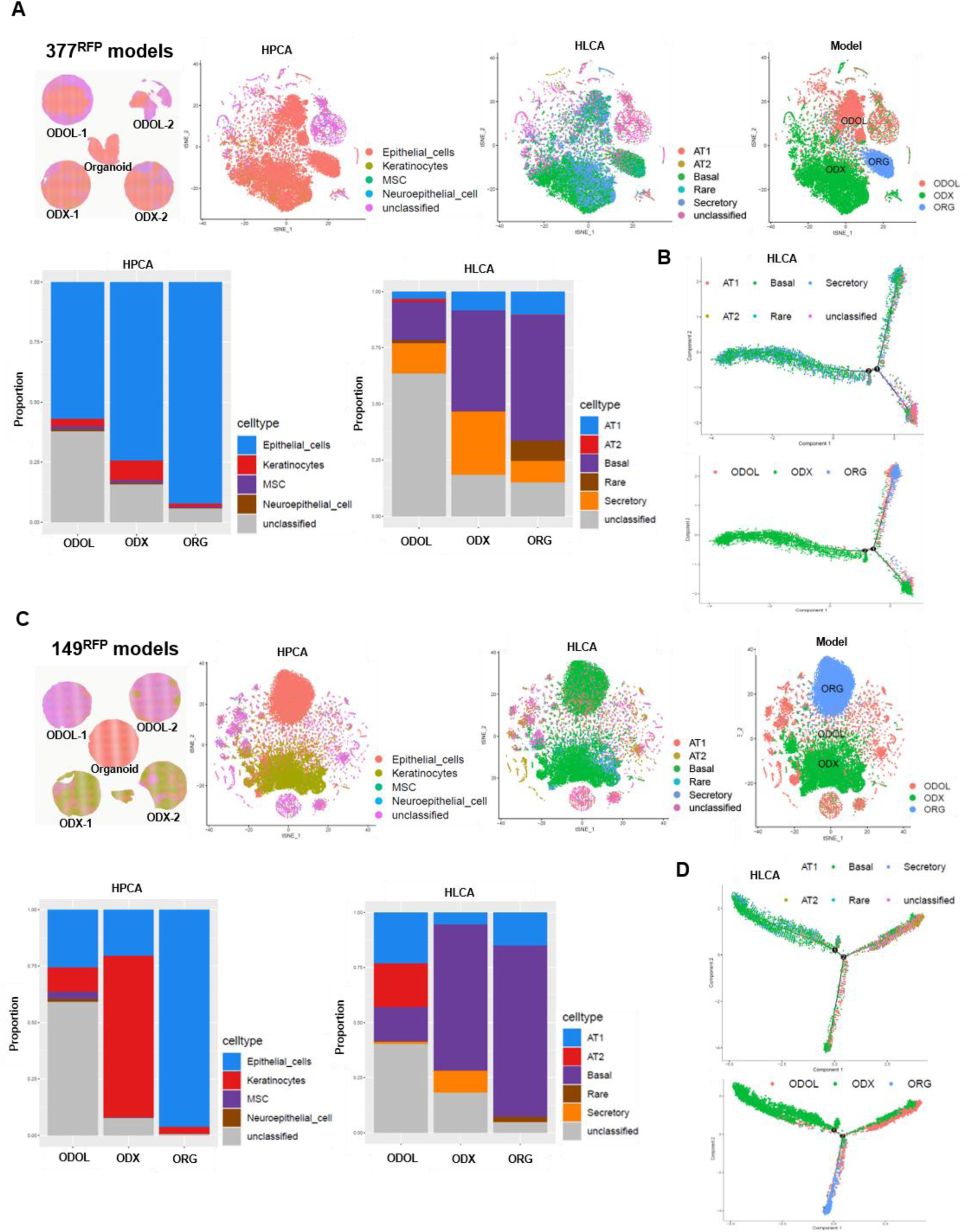
Spatial transcriptomics. (A) tSNE showing cell embeddings among XDO377^RFP^, ODX-377^RFP^, and ODOL-377^RFP^. (B) Pseudotime analysis among XDO377^RFP^, ODX-377^RFP^, and ODOL-377^RFP^. (C) tSNE showing cell embeddings among XDO149^RFP^, ODX-149^RFP^, and ODOL-149^RFP^. (D) Pseudotime analysis among XDO149^RFP^, ODX-149^RFP^, and ODOL-149^RFP^.

## Discussion

We have shown here that intact LUSC organoids have superior ability to grow orthotopically in mouse lungs than dissociated organoid cells. While ODOL tumors manifested poorer squamous lineage differentiation than the subcutaneously growing ODX tumors, they demonstrated slower growth rates *in vivo*, which is linked to the less hypoxic environment of the lung TME. We further showed that the hypoxic condition of subcutaneous TME induced HIF1α, which upregulated SOX2 that is required for maintenance of LUSC lineage and promote differentiation. Paradoxically, the less hypoxic condition of the lung TME induces EMT lineage plasticity and ECM remodeling. Collectively, our study has shown that SOX2 is a key promoter of lineage survival and proliferation, and HIF1α is a key regulator of lineage plasticity and ECM remodeling in LUSC. These findings provide new insights on the complex relationship of TME, hypoxia and SOX2 in shaping lineage plasticity in LUSC (**Figure 8**).

**Figure 8.**
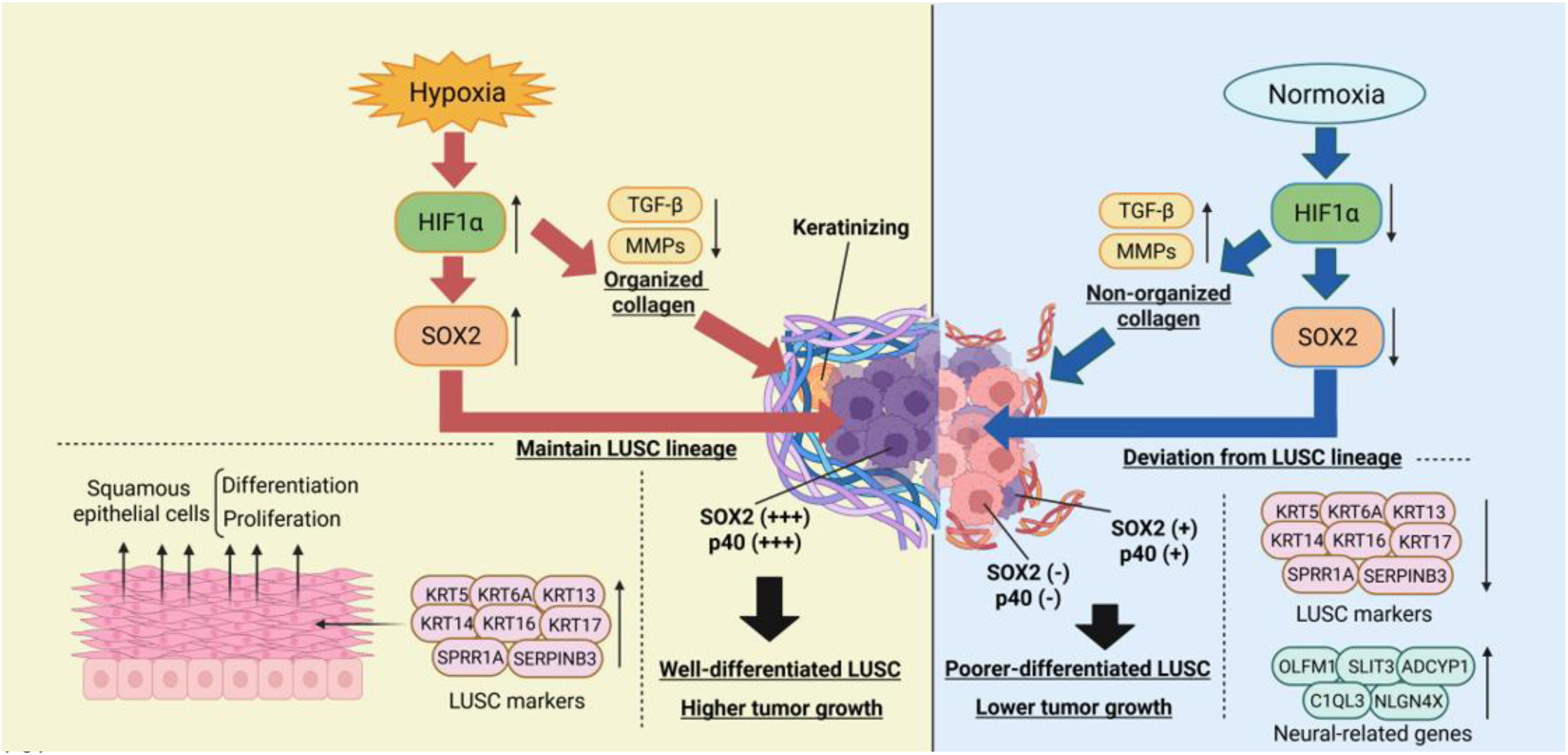
Schematic illustrating the relationship between HIF1α, SOX2, and LUSC lineage. Under hypoxic conditions, HIF1α is activated, while HIF1α simultaneously upregulates SOX2 expression. SOX2 functions as a lineage-survival oncogene in LUSC and contributes to the maintenance of the LUSC lineage by promoting the proliferation and differentiation of squamous epithelium. Furthermore, HIF1α downregulates the expression of TGF-β and MMPs, which are involved in collagen organization. Collectively, these findings indicate that hypoxia promotes LUSC lineage maintenance and tumor progression. In contrast, under non-hypoxic conditions, HIF1α activity is suppressed, and the expression of squamous epithelium-related genes is reduced compared with hypoxia. In addition, suppression of HIF1α upregulates the expression of TGF-β and MMPs, which are involved in non-organized collagen. Moreover, the expression of neural-related genes such as OLFM1, SLIT3, ADCYP1, C1QL3, and NLGN4X was increased in ODOL. These findings indicate that a non-hypoxic condition suppressed the LUSC lineage.

HIF1α has been reported to regulate SOX2 expression in glioblastoma and prostate cancer ^15,16^. The glioblastoma cell lines cultured under hypoxia showed higher expression of HIF1α and SOX2 compared to those cultured under normoxia. Furthermore, knockout of HIF1α led to the suppression of SOX2 under hypoxia. Similarly, in prostate cancer cell lines, SOX2 was found to be upregulated under hypoxia. Additionally, HIF1α knockdown resulted in the suppression of SOX2 under hypoxia. Our study found that when HIF1α was suppressed by SYP-5 or by knock-out using CRISPR/Cas9, SOX2 was suppressed under hypoxia. In contrast, when SOX2 was knocked-out, HIF1α was still upregulated by hypoxia. These findings confirms that HIF1α functions upstream of SOX2 and regulates SOX2 expression in LUSC. Although loss of HIF1α did not completely eliminate SOX2 expression, HIF1α appears to be an important regulator of lineage plasticity in LUSC.

In lung cancer, it has been reported that HIF1α maintains epithelial integrity by repressing TGF-β signaling ^17^. In this case, loss of HIF1α triggers epithelial gene depletion and EMT induction. Our RNA-seq demonstrated reduced keratinization and increased TGF-β-induced protein and EMT in sg*HIF1α*-ODXs. Although EMT typically promotes matrix production, the concurrent upregulation of MMPs in sg*HIF1α*-ODXs to rapid collagen degradation, preventing functional matrix assembly. In this study, MMP14 which directly cleaves collagen types I, II, and III was upregulated in sg*HIF1α*-ODXs, suggesting that MMP14 may primarily contribute to collagen degradation in these models ^18^.

The oxygen concentration in the alveoli is approximately 13.5%, making it one of the most oxygen rich regions in the body ^19,20^. In contrast, peripheral tissues such as subcutaneous tissue have an oxygen concentration of around 5%, a level often referred to as physoxia ^19,20^. Therefore, we set 5% oxygen as the hypoxic condition for organoid culture.

However, the oxygen concentration of lung tumors themselves has been reported to be around 2%, and the expression of *HIF* genes is known to be influenced by both oxygen levels and exposure duration. Thus, further studies are needed to explore even more hypoxic conditions, as well as to investigate the relationship between LUSC lineage and other *HIF* genes such as *HIF1β* and *HIF2α*.

It has been found that the high expression of SOX2 is a critical mechanism for the development and differentiation of a large subset of LUSC, and further, SOX2 expression shows a positive correlation with LUSC differentiation markers ^21–26^. TP63, which encodes p63, and KRT6A, which encodes cytokeratin 6A, were most strongly correlated with SOX2 expression in LUSC ^22^. Furthermore, ectopic expression of SOX2 in the LUAD cell line induced the expression of TP63 and KRT6A ^22^. In our RNA-seq and spatial transcriptomics, LUSC differentiation markers such as KRT6A, KRT14, KRT16, and SERPINB3 were found to be elevated in the SOX2 protein high-expression ODX models. However, while the RNA and protein expression level of SOX2 were consistent in organoids, they did not align in ODXs (**Figure 2E**) (**Supplementary Figure 1, 12C, and 12F)**. This discrepancy requires further investigation.

Delta Np63 expression was found to increase under hypoxia when HIF1α was upregulated. Conversely, suppression of HIF1α prevented the upregulation of Delta Np63. Furthermore, reduced SOX2 also prevented the upregulation of Delta Np63. Delta Np63 is synonymous with p40 and represents the major isoform of p63 ^27^. SOX2 and p63 co-occupy chromatin in LUSC, and chromatin immunoprecipitation sequencing have confirmed their cooperative action at multiple genomic loci ^28,29^. Over 90% of SOX2-positive LUSC cases has been reported to also express p63 ^22,28^. However, in sg*HIF1α*-ODXs and sg*SOX2*-ODXs, the expression of Delta Np63 did not differ significantly from controls. This suggests that, in ODXs, factors other than HIF1α and SOX2 play a major role in regulating Delta Np63, warranting further investigation.

In the DEGs analysis comparing ODX-377^RFP^ and ODOL-377^RFP^, 2,406 genes showed differential expression. However, when organoids derived from each site were compared, most of these differences in gene expression disappeared. These findings indicate that the phenotypes of organoids are modulated by the differences in the TME, and that these phenotypic changes are reversible, reflecting cancer lineage plasticity. Spatial transcriptomics revealed that the phenotypic changes observed in this study are regulated by the lineage plasticity between basal cell-like and AT1/AT2 cell-like tumor cells. Consistent with this, single-cell RNA-seq analyses of patient samples have shown that LUSC is predominantly basal cell-like, whereas LUAD is largely AT2 cell-like ^30^.

Recent studies revealed that lineage plasticity is associated with tumor heterogeneity, histological transformation, and therapeutic resistance in lung cancer, highlighting the growing importance of research in this area ^31–35^. While the understanding of key genetic events associated with lineage plasticity such as SOX2 in LUSC and NKX2-1 in LUAD has advanced, the role of metabolic reprogramming in regulating lineage plasticity remains limited. This is partly due to the difficulty of obtaining both early stage preinvasive and advanced tumor samples from the same patient, the limited heterogeneity of conventional cell lines, and the inability of subcutaneous tumor models to faithfully recapitulate the TME. Using an ODOL model that preserves tumor heterogeneity and the intrapulmonary TME, we demonstrated that hypoxia regulates lineage plasticity in LUSC. This study provides new insights not only for the development of novel therapies targeting the TME but also into histological transformations and their precursor changes associated with resistance to chemotherapy and molecular targeted therapies ^31,32^, potentially contributing to the development of strategies to overcome such resistance.

Nevertheless, hypoxia is only one of several factors that regulate the expression of SOX2. For example, the deubiquitinating enzyme Ubiquitin-Specific Peptidase 13 (USP13) has been reported to modulate SOX2 expression via c-Myc, thereby contributing to the differentiation of LUSC ^36^. To gain a deeper understanding of the developmental process of LUSC, additional studies are necessary for the metabolic reprogramming involved in the regulation of lineage-associated oncogenes.

In conclusions, we established a method for generating an ODOL model and demonstrated that it exhibited phenotypic changes compared to the ODX model. Through analyses using RNA-seq, spatial transcriptomics, and CRISPR/Cas9-based knockout organoid models, we demonstrated that hypoxia, a key component of the TME, regulates this lineage plasticity via the HIF1α-SOX2 axis and ECM remodeling. These results not only enhance our understanding of LUSC biology but also provide important insights for developing novel therapeutic strategies.

## Methods

### Tumor tissue processing, organoid culture and hypoxic condition

The protocols for establishment and characterization of the PDXs and XDOs have been previously described ^6,37^. XDO377 and XDO149 were established from LUSC-PDX models PHLC377-X and PHLC149-X. XDOs were cultured in the M27 medium (**Key resource table 1**) at 37°C with 5% CO2 condition. XDOs were regularly subjected to short tandem repeat analysis and compared with the original tumor to ensure ancestry fidelity and maintained a murine-pathogen-free and mycoplasma-free status. For the hypoxic culture condition, XDOs were incubated for 5 days in a modular incubator chamber (CellXpert C170i, Eppendorf) flushed with a gas mixture containing 5% O_2_ balanced with 5% CO_2_ and N_2_ at 37°C.

### RFP and Venus-Akaluc transduction assay for XDOs

We used lentiviral infection strategy to transduce XDOs. Lentiviruses were prepared as described ^38,39^ by using the plasmids: (i) pMDLg/pRRE, (ii) the vesicular stomatitis virus (VSV-G) envelope plasmid pCMV-VSG, (iii) the rev-expressing plasmid pRSV-Rev, and (iv) the gene transfer vector, a lentiviral vector, TRC-007 turbo red (Turbo RFP) plasmid which is a control plasmid provided by the RNAi Consortium (TRC), or Venus-Akaluc (#124701, Addgene). The infection protocol for XDOs was adapted from Koo, et al ^40^. The XDOs were dissociated to single-cells and these were centrifuged at 800 g for 1 hour at 32°C with M27 media, polybrene (Sigma), and the viral particles, followed by incubation for 4-6 hours.

Subsequently, cells were embedded in Matrigel. RFP in XDOs was confirmed by confocal microscopy (Zeiss LSM700; Zeiss), and RFP in ODOLs and ODXs were confirmed using macroscopy (Zeiss Axiozoom Macroscope; Zeiss). Akaluc signal was confirmed using IVIS Xenogen after reacting the TokeOni (Tocris Bioscience).

### Gene knockout assay for XDOs using CRISPR/Cas9

The CRISPR plasmid (#52961, Addgene) was digested and dephosphorylated with BsmBI (Thermo Scientific) for 30 min at 37 °C using FastAP (Thermo Scientific), 10x FastDigest Buffer (Thermo Scientific), and 100mM DTT (Thermo Scientific). Each pair of oligonucleotides was phosphorylated and annealed using 10x T4 DNA Ligase Buffer (NEB) and T4 PNK (NEB) under the following temperature and cycling conditions: 37 °C for 30 min, 95 °C for 5 min, followed by a gradual cooling to 25 °C at a rate of 5 °C per minute, to generate double-stranded fragments suitable for ligation into the BsmBI-digested CRISPR plasmid, which subsequently induced the expression of single guide RNAs (sgRNAs). The ligation reaction was set up using 2x Quick Ligation Reaction Buffer (NEB) and Quick Ligase (NEB). The resulting product was then transformed into Stbl3 competent cells (Invitrogen).

The plasmid was cultured in Luria-Bertani medium supplemented with ampicillin (Sigma), and DNA was extracted using Plasmid Maxi Kits (QIAGEN). The sgRNA sequences are listed in **Key resource table 2**.

Transduction was performed using electroporation (Neon NxT, Thermo Scientific). The gene/protein knockout was confirmed by targeted DNA sequence and by Western blotting (WB). sg*LacZ* knockout models were used as controls.

### Cell proliferation assay

The XDOs were dissociated into single-cells. Subsequently, 3,000 cells per well were suspended in 75 µL of medium containing 2% Matrigel and seeded onto a base layer of 8 µL Matrigel using a 384-well plate. Cell viability was assessed using CellTiter-Glo 3D (Promega) and quantified with Gen5 (Agilent).

### Orthotopic lung inoculation of XDOs

XDO377 was transduced with lentivirus harboring RFP or Venus-Akaluc to facilitate the identification of tumors within the mouse lungs. After removing the culture media, the organoids were mixed with Cell Recovery Solution (Corning) and incubated on ice for 1 hour to isolate them from the Matrigel. XDO377 was counted, and the cells were divided into four groups: (i) 2 million cells with preserved organoid structures, (ii) 1 million cells with preserved organoid structures, (iii) 2 million cells dissociated into single-cell, and (iv) 1 million cells dissociated into single-cell. The cells were mixed with 25 µL of Matrigel and 25 µL of PBS (WISENT). XDO377 cells were injected intratracheally into 6- to 8-week-old non-obese severe combined immunodeficient gamma (NSG) mice under the surgical microscope, using the method we previously reported with cell lines ^41^. Transplanted mice were followed with 4-weekly computed tomography scan (Mediso nanoScan SPECT/CT/PET) for up to 16 weeks. Prior to scanning, the mice were injected intraperitoneally with the reaction substrate (TokeOni), and Akaluc signal intensity was measured using the bioluminescent system (Xenogen IVIS Spectrum). Following the initial optimization, all subsequent experiments were conducted using intact organoids with 2 million cells. For ODXs, 2 million intact organoids were implanted into the subcutaneous flank of 6- to 8-week-old NSG mice, and tumor growth was monitored using caliper once or twice per week.

### Histology slide preparation and quantitative assessment

Fresh tissue was fixed in 10% formalin for 24-48 hours, transferred to 70% ethanol, and then embedded in a paraffin block. XDOs were recovered using Cell Recovery Solution, suspended in HistoGel (Thermo Scientific), fixed in 10% formalin for 24-48 hours, transferred to 70% ethanol, and then embedded in a paraffin block. Paraffin blocks were cut into 4 µm thick sections and dried overnight. Tissue sections were stained with hematoxylin and eosin (H&E) and immunohistochemistry (IHC) using BenchMark XT autostainer (Ventana Medical Systems). The antibodies used in IHC are listed in **Key resource table 3**. H-score and percentage of positive stained cells were quantitively evaluated using HALO software (Indica Labs).

### Immunofluorescence (IF) staining

Paraffin blocks were cut into 4 µm thick sections and dried overnight at 60°C.

Deparaffinization and hydration were performed using xylene and descending grades of ethanol. Antigen retrieval was performed using 10% Dako Target Retrieval Solution pH9 (Agilent) at 96°C. Sections were permeabilized using 0.1% of Triton X-100 (Sigma) in PBS (PBS-T), and then were blocked in 3% bovine serum albumin (BSA) (WISENT) in PBS-T. The antibodies used in IF are listed in **Key resource table 3**. One µg/mL DAPI (Bio-Rad) was added, and images were captured using confocal microscopy (STELLARIS Confocal Microscope; Leica Microsystems).

### Picrosirius red staining and Second harmonic generation (SHG)

To evaluate the quantity and nature of collagen in the tumor tissue, picrosirius red staining and SHG were performed. Bright-field and polarized images were captured using a Zeiss Axio Imager (Zeiss). Unstained, formalin-fixed, and deparaffinized 10 µm tissue sections were imaged using SHG confocal microscopy (Zeiss LSM 710 NLO two-photon microscope). To evaluate collagen fiber orientation and alignment, the ImageJ plug-in OrientationJ was used. The degree of alignment was assessed from the shape of the orientation distribution, where broad distributions indicated low alignment and sharp peaks indicated high alignment. The degree of alignment was quantified by calculating the ratio between the peak and baseline of each distribution curve.

### DNA extraction, Sanger Sequencing, Whole exome and shallow genome sequencing (WES/sWGS)

XDO DNA was extracted using gSYNCTM DNA Extraction Kit (Geneaid). The quality of DNA was checked by NanoDrop (Thermo Scientific). For PCR, primers, DNA polymerase (AmpliTaq Gold 360, Applied Biosystems), and DNA were mixed, and followed by cycling conditions; After incubation at 95 °C for 10 minutes, 35 cycles of 95 °C for 30 seconds, 54 °C for 30 seconds, and 72 °C for 50 seconds were performed, followed by a final extension at 72 °C for 7 minutes. A gel containing 1-2% agarose in TAE buffer was prepared, and the PCR product was verified by electrophoresis. Impurities were removed using ExoSAP (Applied Biosystems), and the purified product was submitted for Sanger sequencing. PCR primers for Sanger sequencing are listed in **Key resource table 2**.

Genomic DNA was captured using the Agilent SureSelect Human All Exon v5 Capture Kit (Agilent) and sequenced on the Illumina HiSeq 2500 (Illumina) in a 150 bp paired-end run with ∼400x mean coverage. Potential mouse reads were filtered out using Xenome ^42^, then aligned to GRCh37 genome build using Burrows-Wheeler Aligner ^43^, and further quality control, base quality score recalibration and duplicate read removal were performed using the Genome Analysis Toolkit (Broad Institute) and Picard v1.140 ^44^. Algorithms of MuTect2 and VarScan 2.4.3 were used for single-nucleotide and indel variants calling ^45,46^. Databases of dbSNP ^47^, Exome Aggregation Consortium (ExAC) ^48^, exome sequencing project (ESP) ^49^ and gnomAD v2.1.1 ^50^ were used as public references to filter out germline variants. The final mutations were annotated by Annovar ^51^ and Variant Effect Predictor v112 ^52^. The exonic mutations overlapping rates were calculated as the concordance between ODOL, XDO and ODX models.

Shallow whole-genome sequencing (sWGS) was prepared with Agilent SureSelect XT HS2 Library and performed on NovaSeq X Sequencing systems (Illumina) at low coverage (∼1X).The fastq reads were aligned to GRCh37 genome build using Burrows-Wheeler Aligner ^43^, then read counts profile (bin size ∼50 kb) normalization, GC-content correction and copy number alterations was analyzed by ControlFreec ^53^ and ichorCNA ^54^. The copy number correlations were calculated as the concordance between ODOL, XDO and ODX models.

The subclonal deconvolution was also performed between ODOL, XDO and ODX. Briefly, the mutations and CNA were as the input to estimate the cancer cell fractions (CCF) of each mutation, subclonal clustering by Pylone ^55^. The citup was then used to construct the clonal trees ^56^.

### RNA extraction and RNA-seq

Tumors from xenografts were homogenized in TRIzol (Ambion), and RNA was isolated and precipitated with chloroform and 70% ethanol. XDOs were recovered using Cell Recovery Solution. Total RNA was extracted using the RNeasy Mini Kit (QIAGEN). RNA was also extracted from FFPE blocks using the RNeasy FFPE Kit (QIAGEN). DNase treatment is performed using 10x DNase I Buffer (Invitrogen), DNase I (Invitrogen), and DNase Inactivation Reagent (Invitrogen). The quality of RNA was checked by NanoDrop, Qubit RNA HS Assay kit (Thermo Scientific), and Bioanalyzer (Agilent).

Library preparation was performed using the Illumina Stranded mRNA Prep Ligation Kit (Illumina) with 200 ng of RNA as input. Libraries were sequenced on a NovaSeq X system using 150-cycle paired-end reads. Xenome was used to filter mouse reads from human reads ^42^. The reads then were aligned by STAR aligner ^57^, RSEM were used to obtain gene-level counts and transcripts per million (TPM) values from the resulting BAM files ^58^. ComBat was applied to adjust for batch effects ^59^. Top 2000 most variable genes were selected for the PCA. The DEGs were obtained by DESeq2 in R ^60^. GSEA was performed with the GSEA (Broad Institute, version: 4.0.3) using MSigDB Hallmark and REACTOME database ^61,62^.

### Reverse-transcription quantitative polymerase chain reaction (RT-qPCR)

RNA was reverse transcribed to cDNA using Oligo dT Primer (Thermo Scientific), dNTP Mix (Fermentas), 5x First Stand Buffer (Invitrogen), 0.1M DTT (Invitrogen), and SuperScript III (Invitrogen). qPCR was performed using the SYBR Green method (Applied Biosystems), with the following conditions; 95°C for 30 seconds, 60°C for 1 minute, 95°C for 1 minute, 60°C for 30 seconds, and 95°C for 30 seconds for 40 cycles. The primers for RT-qPCR are listed in **Key resource table 2**.

### Western blotting (WB)

XDOs were lysed with RIPA Buffer (Sigma-Aldrich) with Protease inhibitor cocktail tablets (Roche) and Phosphatase inhibitor mini tablets (Thermo Scientific). The extracted proteins were mixed with the BCA protein assay kit (Thermo Scientific), and then quantified using Gen5. Proteins were denatured in sample buffer (Bio-Rad), and loaded for SDS-PAGE (Bio-Rad). Proteins were transferred onto nitrocellulose membranes (Bio-Rad) and blocked in 5% skim milk or 5% BSA, and probed with antibodies. The antibodies are listed in **Key resource table 3**. ECL (Bio-Rad) reagents were used for detecting protein bands.

### Spatial transcriptomics (10x Visium HD)

FFPE blocks were cored using a 15 mm punch, and a tissue microarray was constructed. Five-micrometer-thick tissue sections were prepared from tissue microarray and mounted on Visium Spatial Gene Expression slides (10x Genomics) according to the manufacturer’s instructions. Tissue sections were deparaffinized, fixed, and H&E stained for histological visualization. Imaging was performed using a high-resolution brightfield microscope. Spatially barcoded mRNA capture was performed using the Visium HD Gene Expression workflow. Libraries were generated following the 10x Genomics protocol and sequenced on an Illumina NovaSeq 6000 platform to a targeted depth of approximately 50,000 reads per capture spot. Space Ranger (v.4.0, 10x Genomics) results were used to perform expression analysis, mapping, counting and clustering. The reads count matrix were loaded into Seurat v5.3 for all subsequent data quality control, filtering, normalization, dimensional reduction and visualization ^63^. The k-nearest neighbours FindNeighbors and FindClusters functions in Seurat were performed to get the clusters. The deconvolution of cells in each spot was applied to get the compositions using the RCTD algorithm based on the annotated scRNA-seq data of Human Lung Cell Atlas (HLCA) ^64^. We then annotated the cells in the spot using singleR with Human Primary Cell Atlas (HPCA) dataset ^65^. To reconstruct the spatial trajectory in organoid, ODX and ODOL models, we selected the singlet and epithelial cells in the spot from RCTD deconvolution output, Monocle then was used to infer the pseudotime and temporal cell trajectories ^66^.

## Statistical analysis

Data are presented as the means ± standard deviation (SD), and GraphPad Prizm version 10.4.0 (GraphPad Prism Corp.) was used for statistical analyses. Two-way ANOVA was used to examine the differences between and within groups in repeated measurement data. Significant differences were estimated using Welch’s t test. The level of significance is indicated by the P value in each experiment. Asterisks in the figures indicate the following: *P < 0.05; **P < 0.01; ***P < 0.001; ****P < 0.0001; ns. P > 0.05.

**Key resource table 1.** Lung cancer organoid culture media (M27)

| Materials | Final concentration | Stock concentration | 10mL of media |
| --- | --- | --- | --- |
| Advanced DMEM/F-12 (Gibco) |  |  | 10 mL |
| B-27 Supplement (Gibco) | 1x | 50x | 200 µL |
| GultaMAX (Gibco) | 2 mM | 200 mM | 100 µL |
| HEPES (Gibco) | 10 mM | 1 M | 100 µL |
| Antibiotic-Antimycotic (Gibco) | 100 U/mL | 10000 U/mL | 100 µL |
| N-Acetyl-L-cysteine (Sigma) | 1.25 mM | 500 mM | 25 µL |
| Recombinant Human FGF-4 (Peprotech) | 100 ng/mL | 100 µg/mL | 10 µL |
| Recombinant Human FGF-7 (Peprotech) | 5 ng/mL | 100 µg/mL | 0.5 µL |
| Recombinant Human EGF (Corning) | 50 ng/mL | 100 µg/mL | 5 µL |
| Recombinant Human Noggin (Peprotech) | 100 ng/mL | 100 µg/mL | 10 µL |
| A83-01 (Tocris) | 0.5 µM | 0.5 mM | 10 µL |
| Y-27632 (Selleck Chemicals) | 10 µM | 10 mM | 10 µL |
| SAG (Enzo) | 100 nM | 1 mM | 1 µL |
| CHIR 99021 (Tocris) | 250 nM | 5 mM | 0.5 µL |

**Key resource table 2.**
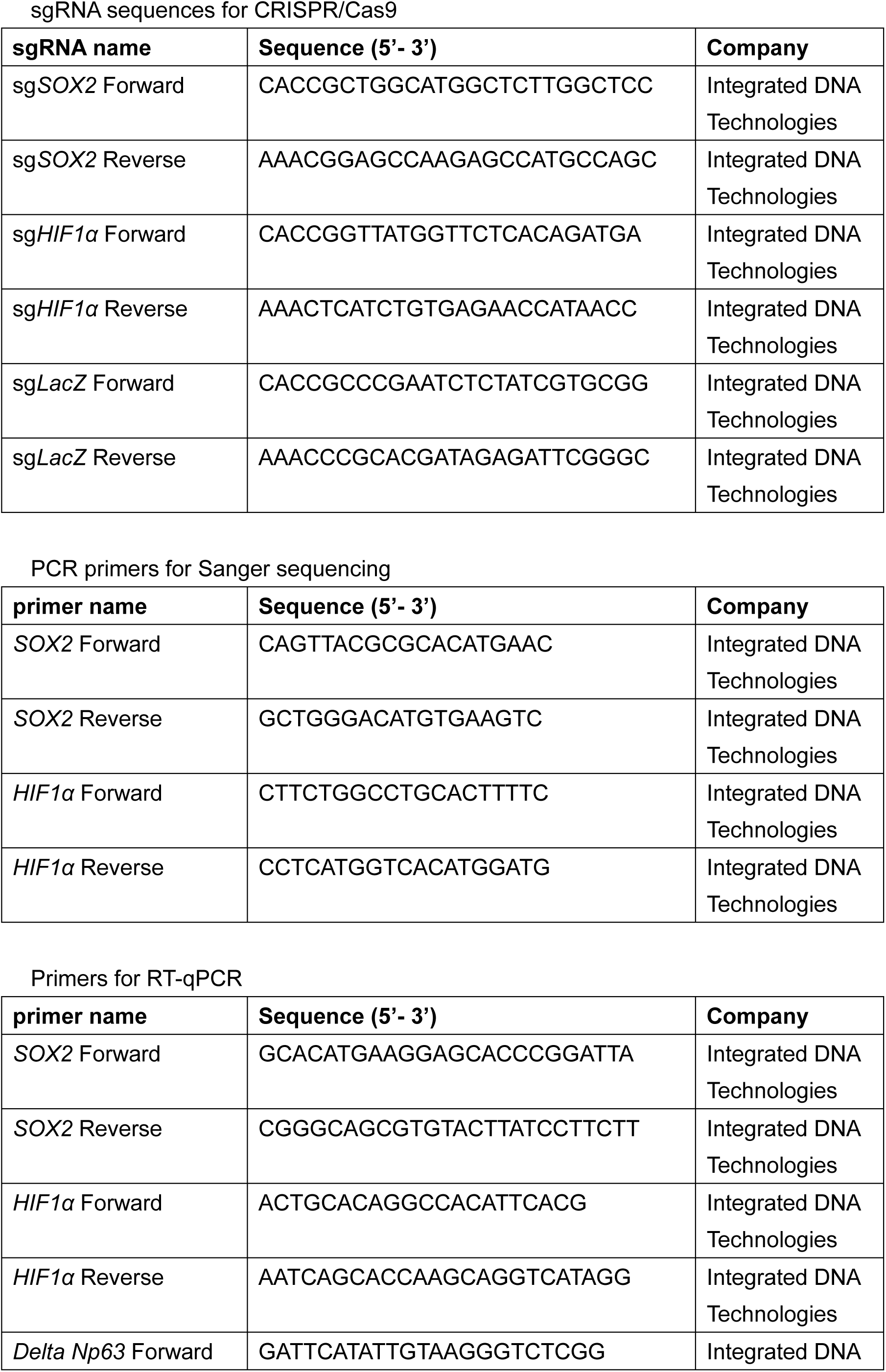
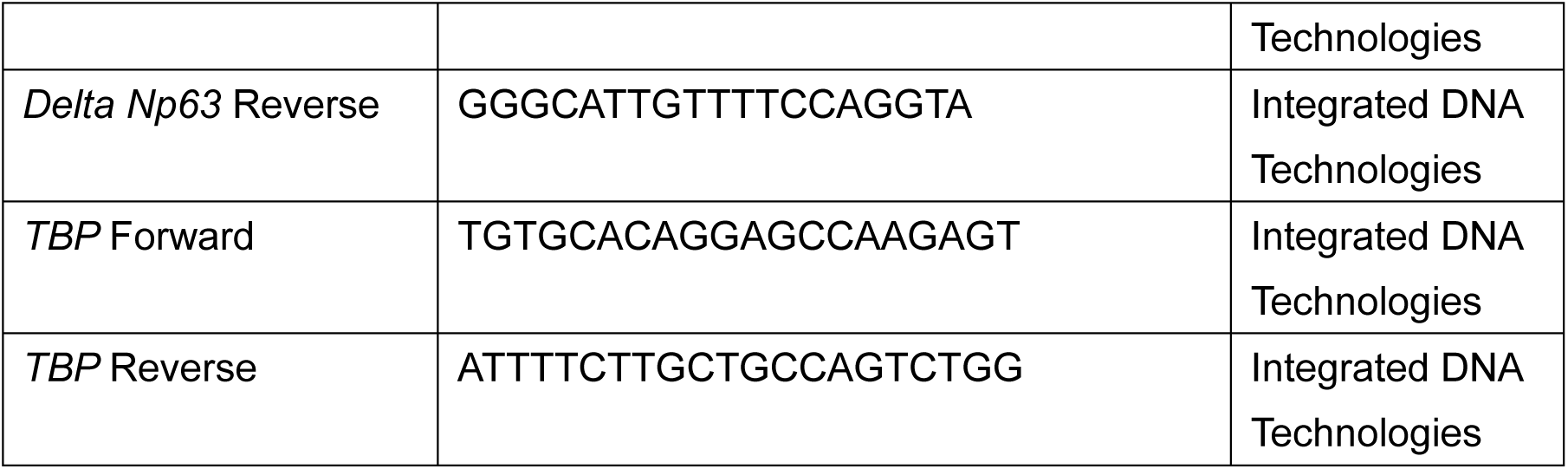
List of primers.

**Key resource table 3.** List of antibodies.

| Antibodies name | Clone | Concentration | Application | Company |
| --- | --- | --- | --- | --- |
| p40 | BC28 | 1 drop-on optiview program | IHC | VENTANA |
| SOX2 | AF2018 | 1:4000 | IHC | R&D |
| HIF1α | EP1215Y | 1:100 | IHC | BIOCARE |
| SOX2 | AF2018 | 1:200 | IF | R&D |
| HIF1α | Clone54 | 1:100 | IF | BD Bioscience |
| Alexa Fluor 647 donkey anti-goat | A21447 | 1:200 | IF | Molecular Probes |
| Alexa Fluor 568 rabbit anti-mouse | A11061 | 1:200 | IF | Invitrogen |
| SOX2 | AF2018 | 1:1000 | WB | R&D |
| Delta Np63 | E6Q3O | 1:1000 | WB | Cell Signaling |
| HIF1α | #3716 | 1:1000 | WB | Cell Signaling |
| β-actin | #4967 | 1:1000 | WB | Cell Signaling |
| anti-rabbit IgG HRP-linked antibody | #7074 | 1:3000 | WB | Cell Signaling |
| donkey anti-goat IgG HRP-linked antibody | sc-2033 | 1:3000 | WB | Santa Cruz |

## List of abbreviations

LUSC: lung squamous cell carcinoma
XDO: xenograft-derived organoid
ODOL: organoid-derived orthotopic lung
ODX: organoid-derived xenograft
HIF1α: hypoxia-inducible factor 1-alpha
SOX2: SRY-box transcription factor 2
ECM: extracellular matrix
TME: tumor microenvironment
NSCLC: non-small cell lung cancer
LUAD: lung adenocarcinoma
LCO: lung cancer organoid
PDX: patient-derived xenograft
RFP: red fluorescence protein
sgRNA: single guide RNA
NSG mice: non-obese severe combined immunodeficient gamma mice
H&E: hematoxylin and eosin
IHC: immunohistochemistry
IF: immunofluorescence staining
SHG: second harmonic generation
WGS: whole genome sequencing
sWGS: shallow genome sequencing
RNA-seq: RNA sequencing
RT-qPCR: reverse-transcription quantitative polymerase chain reaction
WB: Western blotting
CCF: cancer cell fractions
TPM: transcripts per million
PCA: principal component analysis
DEGs: differentially expressed genes
GSEA: gene set enrichment analysis
FFPE: formalin-fixed paraffin-embedded
HLCA: Human Lung Cell Atlas
HPCA: Human Primary Cell Atlas
KRT: Keratin
SPRR1A: Small Proline-Rich Protein 1A
SERPINB: Serpin Family B Member
NKX2.1: NK2 Homeobox 1
NLGN4X: Neuroligin 4 X-Linked
C1QL3: Complement C1q Like 3
ADCYAP1: Adenylate Cyclase Activating Polypeptide 1
OLFM1: Olfactomedin 1
BNIP3: BCL2 Interacting Protein 3
PFKFB3: 6-Phosphofructo-2-kinase/fructose-2,6-bisphosphatase 3
FAM162A: Family with Sequence Similarity 162 Member A
mTOR: mechanistic target of rapamycin
EMT: epithelial-mesenchymal transformation
TGFI: transforming growth factor beta induced protein
MMP: matrix metalloproteinases
AT1: alveolar type 1 AT2: alveolar type 2
USP: Ubiquitin-Specific Peptidase

## Acknowledgements

We thank Shavanie Sheecharran and Jian Ying Zhou from the AMPL laboratory at Princess Margaret Cancer Centre for their technical assistance. YF was supported by the Japan Society for the Promotion of Science (JSPS) Overseas Challenge Program for Young Researchers. This work was supported by the Canadian Institutes of Health Research (CIHR) Foundation (grant number: FDN-148395) to MST, the Uehara Memorial Foundation and the Grants-in-Aid for Scientific Research JSPS KAKENHI (Grant number: 24K184850202) to HO, the JSPS KAKENHI (grant number: 20KK0202 and 25K02728) to TA and YM, the William Coco Chair in Surgical Innovation for Lung Cancer to YK, and the Princess Margaret Cancer Foundation.

## Author contributions

Conceptualization, YF, HO, MST, KY ; Methodology, YF, HO, QL, RN, TK, NH, NR, HH, TS, YH, NR, MST; Investigation, YF, HO, QL, FY, TY, and MST; Formal Analysis, YF, HO, QL, MST; Resources, YF, HO, QL, RN, TK, NR, HH, NB, TS, YH, FY, TY, TA, YM, NR, MST, KY; Visualization, YF, HO, QL, RN, TK, NR, HH, NB, TS, YH, FY, TY, TA, YM, MST, KY; Funding Acquisition, YF, HO, TA, YM, MST, KY; Project Administration, MST and KY.

## Data Availability

The data that support the findings of this study are available from the corresponding author upon reasonable request.

## Declarations

### Ethics approval and consent to participate

The collection of surgically resected primary tumors from patients and the development of PDXs were approved by the University Health Network Research Ethics Board (REB: 17-558) and Animal Care Committee (AUP: 5555). Informed written consent was received from all patients. Animal study to establish ODOL and ODX was done under Animal Care Committee protocol (AUP: 6087) from the Institutional Animal Care and Use Committee of the University Health Network.

### Consent for publication

Not applicable

## References

1. Bray F., et al. Global cancer statistics 2022: GLOBOCAN estimates of incidence and mortality worldwide for 36 cancers in 185 countries. CA Cancer J Clin 74, 229–263 (2024).

2. Nicholson A. G., et al. The 2021 WHO Classification of Lung Tumors: Impact of Advances Since 2015. J Thorac Oncol 17, 362–387 (2022).

3. Wang B. Y., et al. The comparison between adenocarcinoma and squamous cell carcinoma in lung cancer patients. J Cancer Res Clin Oncol 146, 43–52 (2020).

4. Hao B., et al. Squamous cell carcinoma predicts worse prognosis than adenocarcinoma in stage IA lung cancer patients: A population-based propensity score matching analysis. Front Surg 9, 944032 (2022).

5. Sachs N., et al. Long-term expanding human airway organoids for disease modeling. Embo j 38, (2019).

6. Shi R., et al. Organoid Cultures as Preclinical Models of Non-Small Cell Lung Cancer. Clin Cancer Res 26, 1162–1174 (2020).

7. Kim M., et al. Patient-derived lung cancer organoids as in vitro cancer models for therapeutic screening. Nat Commun 10, 3991 (2019).

8. Bejarano L., Jordāo M. J. C., Joyce J. A. Therapeutic Targeting of the Tumor Microenvironment. Cancer Discov 11, 933–959 (2021).

9. Kahn J., Tofilon P. J., Camphausen K. Preclinical models in radiation oncology. Radiat Oncol 7, 223 (2012).

10. Masters J. R. Human cancer cell lines: fact and fantasy. Nat Rev Mol Cell Biol 1, 233–236 (2000).

11. Yu K., et al. Comprehensive transcriptomic analysis of cell lines as models of primary tumors across 22 tumor types. Nat Commun 10, 3574 (2019).

12. Fujii M., Sato T. Somatic cell-derived organoids as prototypes of human epithelial tissues and diseases. Nat Mater 20, 156–169 (2021).

13. Kang Y., et al. Proliferation of human lung cancer in an orthotopic transplantation mouse model. Exp Ther Med 1, 471–475 (2010).

14. Junttila M. R., de Sauvage F. J. Influence of tumour micro-environment heterogeneity on therapeutic response. Nature 501, 346–354 (2013).

15. Wang P., et al. HIF1α/HIF2 α -Sox2/Klf4 promotes the malignant progression of glioblastoma via the EGFR-PI3K/AKT signalling pathway with positive feedback under hypoxia. Cell Death Dis 12, 312 (2021).

16. Bae K. M., Dai Y., Vieweg J., Siemann D. W. Hypoxia regulates SOX2 expression to promote prostate cancer cell invasion and sphere formation. Am J Cancer Res 6, 1078–1088 (2016).

17. Ando A., et al. Repressive role of stabilized hypoxia inducible factor 1 α expression on transforming growth factor β -induced extracellular matrix production in lung cancer cells. Cancer Sci 110, 1959–1973 (2019).

18. Hotary K. B., Allen E. D., Brooks P. C., Datta N. S., Long M. W., Weiss S. J. Membrane type I matrix metalloproteinase usurps tumor growth control imposed by the three-dimensional extracellular matrix. Cell 114, 33–45 (2003).

19. McKeown S. R. Defining normoxia, physoxia and hypoxia in tumours-implications for treatment response. Br J Radiol 87, 20130676 (2014).

20. Le Q. T., et al. An evaluation of tumor oxygenation and gene expression in patients with early stage non-small cell lung cancers. Clin Cancer Res 12, 1507–1514 (2006).

21. Kim B. R., et al. SOX2 and PI3K Cooperate to Induce and Stabilize a Squamous-Committed Stem Cell Injury State during Lung Squamous Cell Carcinoma Pathogenesis. PLoS Biol 14, e1002581 (2016).

22. Bass A. J., et al. SOX2 is an amplified lineage-survival oncogene in lung and esophageal squamous cell carcinomas. Nat Genet 41, 1238–1242 (2009).

23. McCaughan F., et al. Progressive 3q amplification consistently targets SOX2 in preinvasive squamous lung cancer. Am J Respir Crit Care Med 182, 83–91 (2010).

24. Maier S., et al. SOX2 amplification is a common event in squamous cell carcinomas of different organ sites. Hum Pathol 42, 1078–1088 (2011).

25. van Boerdonk R. A., et al. DNA copy number alterations in endobronchial squamous metaplastic lesions predict lung cancer. Am J Respir Crit Care Med 184, 948–956 (2011).

26. Schneider F., Luvison A., Cieply K., Dacic S. Sex-determining region Y-box 2 amplification in preneoplastic squamous lesions of the lung. Hum Pathol 44, 706–711 (2013).

27. Bishop J. A., Teruya-Feldstein J., Westra W. H., Pelosi G., Travis W. D., Rekhtman N. p40 (ÄNp63) is superior to p63 for the diagnosis of pulmonary squamous cell carcinoma. Mod Pathol 25, 405–415 (2012).

28. Watanabe H., et al. SOX2 and p63 colocalize at genetic loci in squamous cell carcinomas. J Clin Invest 124, 1636–1645 (2014).

29. Comprehensive genomic characterization of squamous cell lung cancers. Nature 489, 519–525 (2012).

30. Lambrechts D., et al. Phenotype molding of stromal cells in the lung tumor microenvironment. Nat Med 24, 1277–1289 (2018).

31. Quintanal-Villalonga Á., et al. Lineage plasticity in cancer: a shared pathway of therapeutic resistance. Nat Rev Clin Oncol 17, 360–371 (2020).

32. Le Magnen C., Shen M. M., Abate-Shen C. Lineage Plasticity in Cancer Progression and Treatment. Annu Rev Cancer Biol 2, 271–289 (2018).

33. Kaiser A. M., et al. p53 governs an AT1 differentiation programme in lung cancer suppression. Nature 619, 851–859 (2023).

34. Han G., et al. An atlas of epithelial cell states and plasticity in lung adenocarcinoma. Nature 627, 656–663 (2024).

35. Li Z., et al. Alveolar Differentiation Drives Resistance to KRAS Inhibition in Lung Adenocarcinoma. Cancer Discov 14, 308–325 (2024).

36. Kwon J., et al. USP13 drives lung squamous cell carcinoma by switching lung club cell lineage plasticity. Mol Cancer 22, 204 (2023).

37. Mirhadi S., et al. Integrative analysis of non-small cell lung cancer patient-derived xenografts identifies distinct proteotypes associated with patient outcomes. Nat Commun 13, 1811 (2022).

38. Naldini L., et al. In vivo gene delivery and stable transduction of nondividing cells by a lentiviral vector. Science 272, 263–267 (1996).

39. Dull T., et al. A third-generation lentivirus vector with a conditional packaging system. J Virol 72, 8463–8471 (1998).

40. Koo B. K., et al. Controlled gene expression in primary Lgr5 organoid cultures. Nat Methods 9, 81–83 (2011).

41. Nakajima T., et al. Orthotopic lung cancer murine model by nonoperative transbronchial approach. Ann Thorac Surg 97, 1771–1775 (2014).

42. Conway T., et al. Xenome--a tool for classifying reads from xenograft samples. Bioinformatics 28, i172–178 (2012).

43. Li H., Durbin R. Fast and accurate short read alignment with Burrows-Wheeler transform. Bioinformatics 25, 1754–1760 (2009).

44. McKenna A., et al. The Genome Analysis Toolkit: a MapReduce framework for analyzing next-generation DNA sequencing data. Genome Res 20, 1297–1303 (2010).

45. Cibulskis K., et al. Sensitive detection of somatic point mutations in impure and heterogeneous cancer samples. Nat Biotechnol 31, 213–219 (2013).

46. Koboldt D. C., et al. VarScan 2: somatic mutation and copy number alteration discovery in cancer by exome sequencing. Genome Res 22, 568–576 (2012).

47. Sherry S. T., et al. dbSNP: the NCBI database of genetic variation. Nucleic Acids Res 29, 308–311 (2001).

48. Lek M., et al. Analysis of protein-coding genetic variation in 60,706 humans. Nature 536, 285–291 (2016).

49. Fu W., et al. Analysis of 6,515 exomes reveals the recent origin of most human protein-coding variants. Nature 493, 216–220 (2013).

50. Karczewski K. J., et al. The mutational constraint spectrum quantified from variation in 141,456 humans. Nature 581, 434–443 (2020).

51. Yang H., Wang K. Genomic variant annotation and prioritization with ANNOVAR and wANNOVAR. Nat Protoc 10, 1556–1566 (2015).

52. McLaren W., et al. The Ensembl Variant Effect Predictor. Genome Biol 17, 122 (2016).

53. Boeva V., et al. Control-free calling of copy number alterations in deep-sequencing data using GC-content normalization. Bioinformatics 27, 268–269 (2011).

54. Adalsteinsson V. A., et al. Scalable whole-exome sequencing of cell-free DNA reveals high concordance with metastatic tumors. Nat Commun 8, 1324 (2017).

55. Roth A., et al. PyClone: statistical inference of clonal population structure in cancer. Nat Methods 11, 396–398 (2014).

56. Malikic S., McPherson A. W., Donmez N., Sahinalp C. S. Clonality inference in multiple tumor samples using phylogeny. Bioinformatics 31, 1349–1356 (2015).

57. Dobin A., et al. STAR: ultrafast universal RNA-seq aligner. Bioinformatics 29, 15–21 (2013).

58. Li B., Dewey C. N. RSEM: accurate transcript quantification from RNA-Seq data with or without a reference genome. BMC Bioinformatics 12, 323 (2011).

59. Johnson W. E., Li C., Rabinovic A. Adjusting batch effects in microarray expression data using empirical Bayes methods. Biostatistics 8, 118–127 (2007).

60. Love M. I., Huber W., Anders S. Moderated estimation of fold change and dispersion for RNA-seq data with DESeq2. Genome Biol 15, 550 (2014).

61. Subramanian A., et al. Gene set enrichment analysis: a knowledge-based approach for interpreting genome-wide expression profiles. Proc Natl Acad Sci U S A 102, 15545–15550 (2005).

62. Liberzon A., Subramanian A., Pinchback R., Thorvaldsdóttir H., Tamayo P., Mesirov J. P. Molecular signatures database (MSigDB) 3.0. Bioinformatics 27, 1739–1740 (2011).

63. Hao Y., et al. Dictionary learning for integrative, multimodal and scalable single-cell analysis. Nat Biotechnol 42, 293–304 (2024).

64. Cable D. M., et al. Robust decomposition of cell type mixtures in spatial transcriptomics. Nat Biotechnol 40, 517–526 (2022).

65. Sikkema L., et al. An integrated cell atlas of the lung in health and disease. Nat Med 29, 1563–1577 (2023).

66. Qiu X., et al. Reversed graph embedding resolves complex single-cell trajectories. Nat Methods 14, 979–982 (2017).

