## Supplementary Figures for "Tumor Microenvironment Modulates Lineage Plasticity in Lung Squamous Cell Carcinoma"

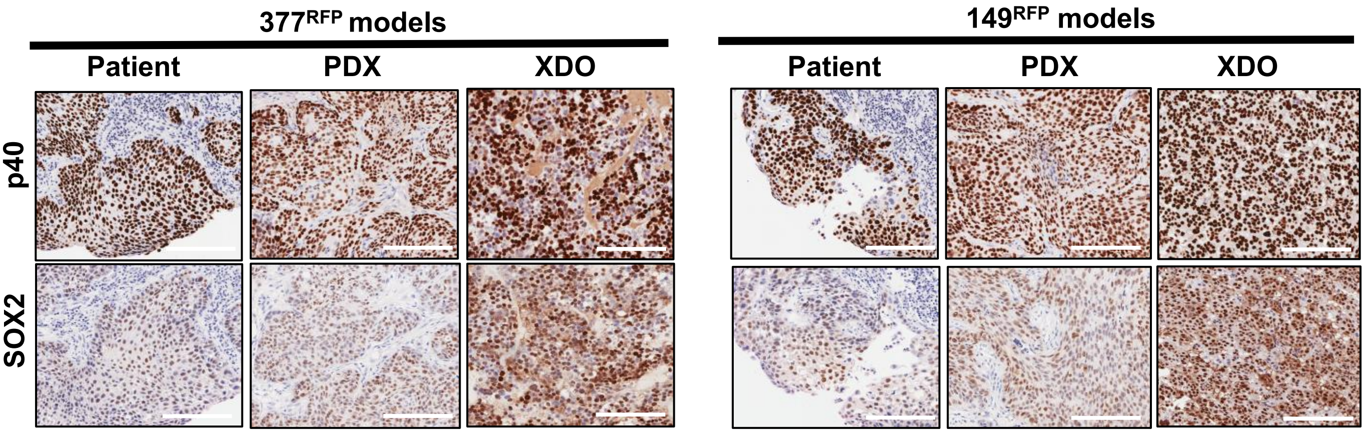


**Supplementary Figure 1.** IHC staining of patient tumors, PDXs, and XDOs in $377^{RFP}$ and $149^{RFP}$ models. These all models showed the positivity for p40 and SOX2. Scale bars are 200 μm.

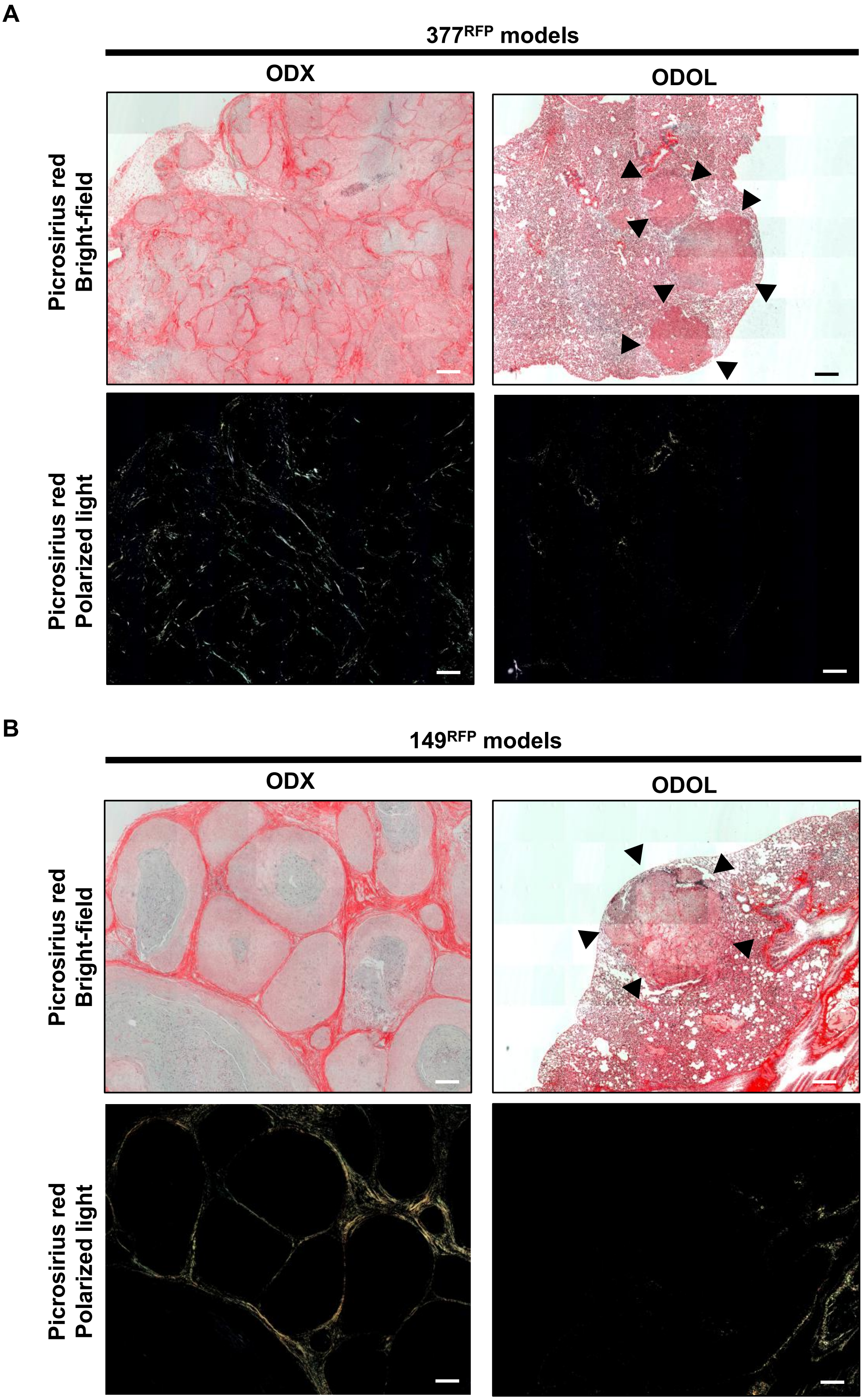


**Supplementary Figure 2.** (A) Picrosirius red staining between ODX-$377^{RFP}$ and ODOL-$377^{RFP}$ (B) Picrosirius red staining between ODX-$149^{RFP}$ and ODOL-$149^{RFP}$. Black arrows indicate tumors. The red color in the bright-field images indicates collagen fibers. The yellow color in the polarized light images indicates collagen type I. The green color in the polarized light images indicates collagen type III. Scale bars are 200 μm.

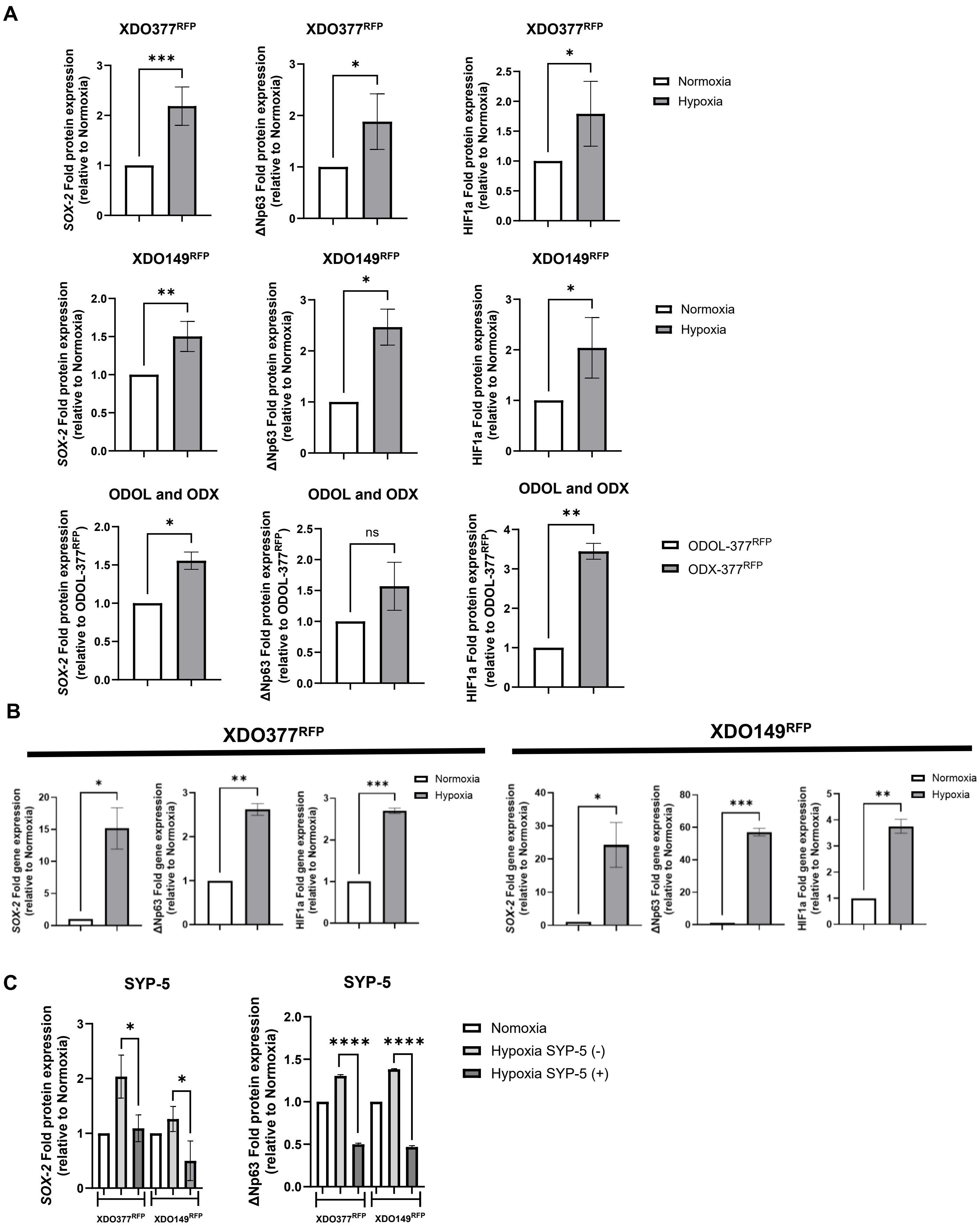


**Supplementary Figure 3.** (A) The quantitative analysis of the western blot in Figure 3D. (B) RT-qPCR between Normoxia and Hypoxia XDOs. (C) The quantitative analysis of the western blot in Figure 3E.

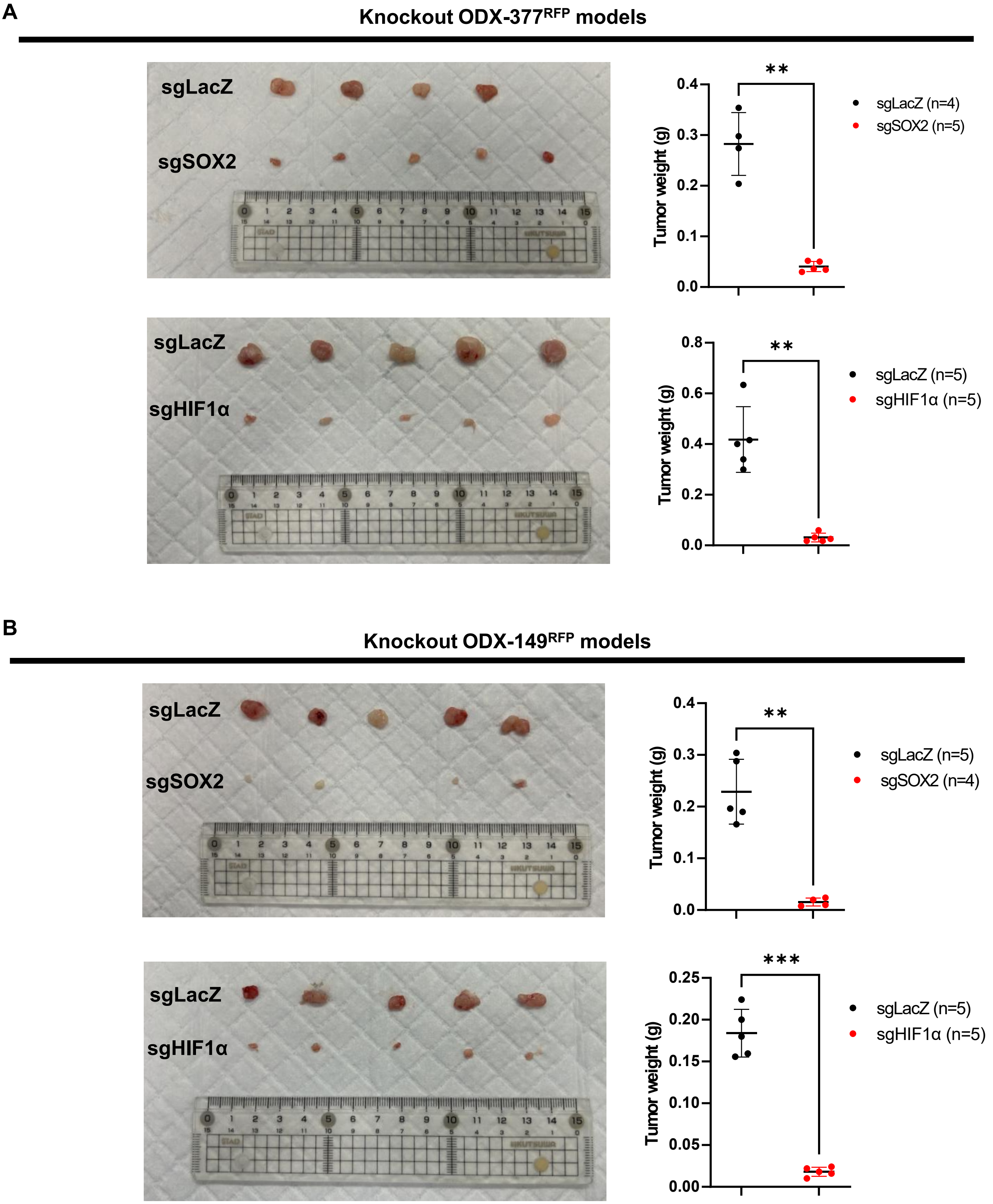


**Supplementary Figure 4.** (A) Tumor pictures and tumor weight of ODX-377$^{RFP\text{-}sgLacZ/\text{-}sgSOX2/\text{-}sgHIF1\alpha}$. (B) Tumor pictures and tumor weight of ODX-149$^{RFP\text{-}sgLacZ/\text{-}sgSOX2/\text{-}sgHIF1\alpha}$.

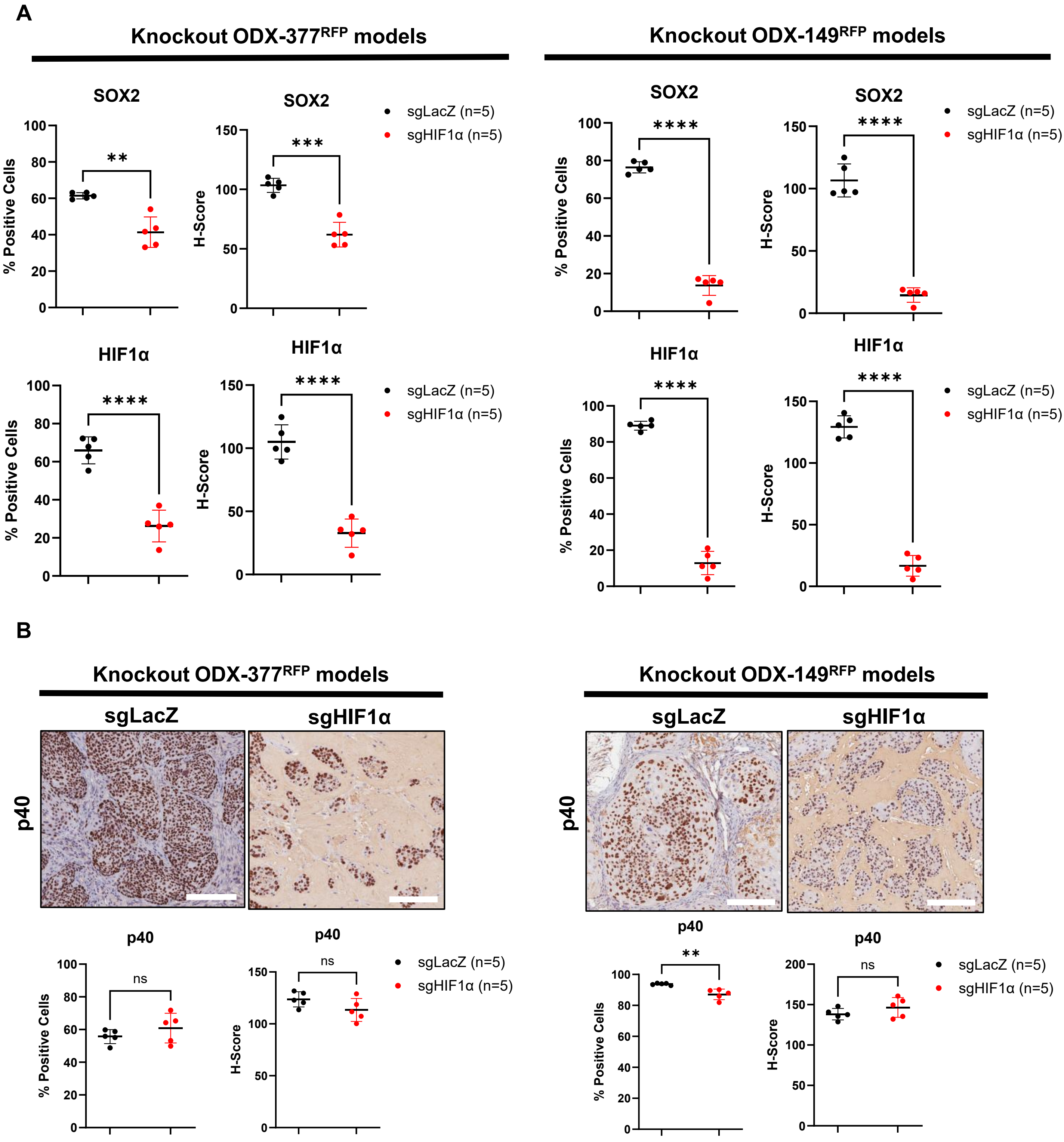


**Supplementary Figure 5.** (A) The staining intensity in Figure 4F was quantitatively evaluated using % positive stained cells and H-score with HALO software. (B) IHC staining for p40 between ODX-377$^{RFP-sgLacZ}$/-149$^{RFP-sgLacZ}$ vs ODX-377$^{RFP-sgHIF1\alpha}$/-149$^{RFP-sgHIF1\alpha}$. Scale bars are 200 μm. The staining intensity was quantitatively evaluated using % positive stained cells and H-score with HALO software.

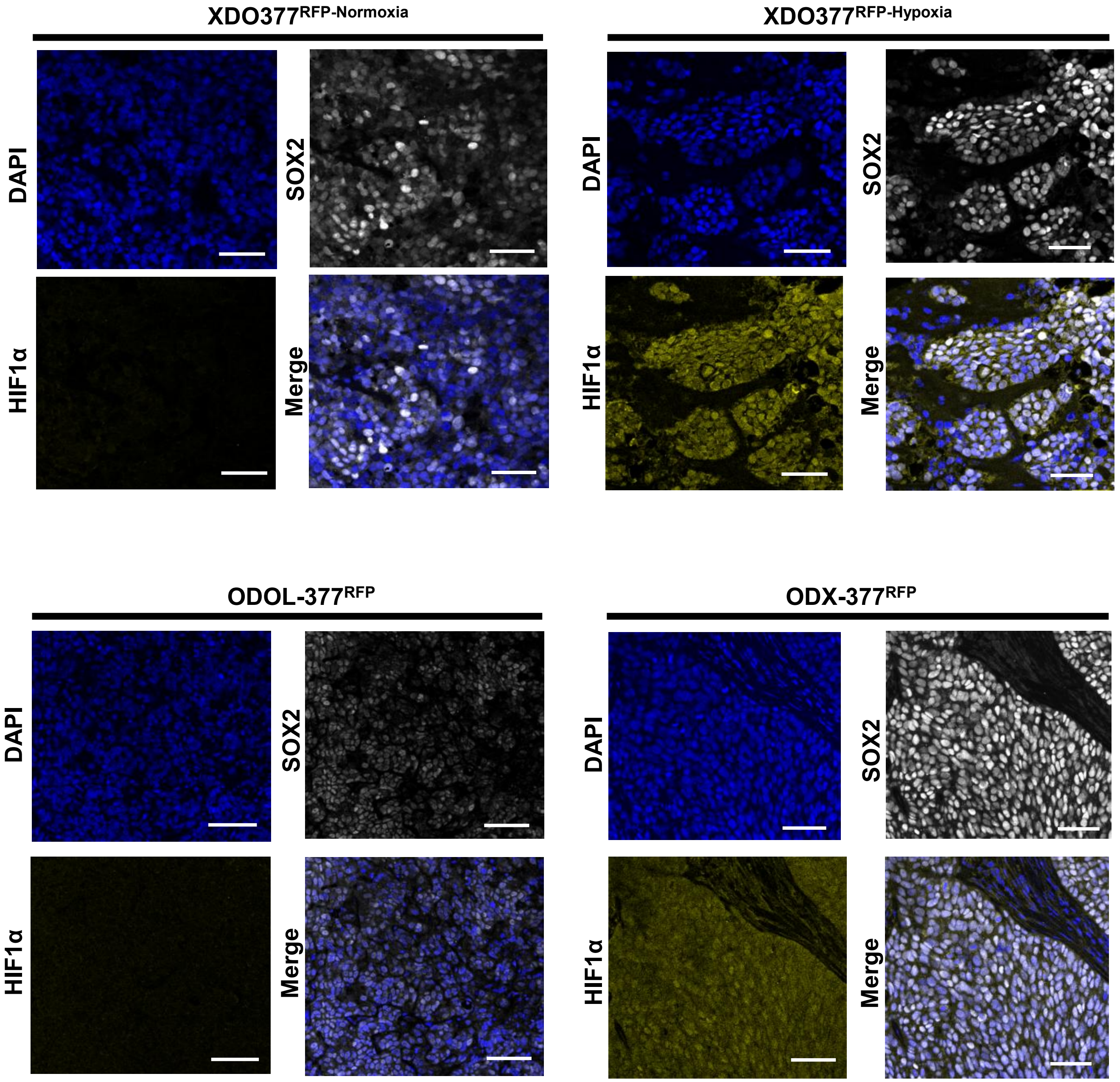


**Supplementary Figure 6.** Immunofluorescence staining in XDO377$^{RFP\text{-}normoxia}$, XDO377$^{RFP\text{-}hypoxia}$, ODX-377$^{RFP}$, and ODOL-377$^{RFP}$ with DAPI, SOX2, and HIF1α. The merge combined these three. Scale bars are 50 μm.

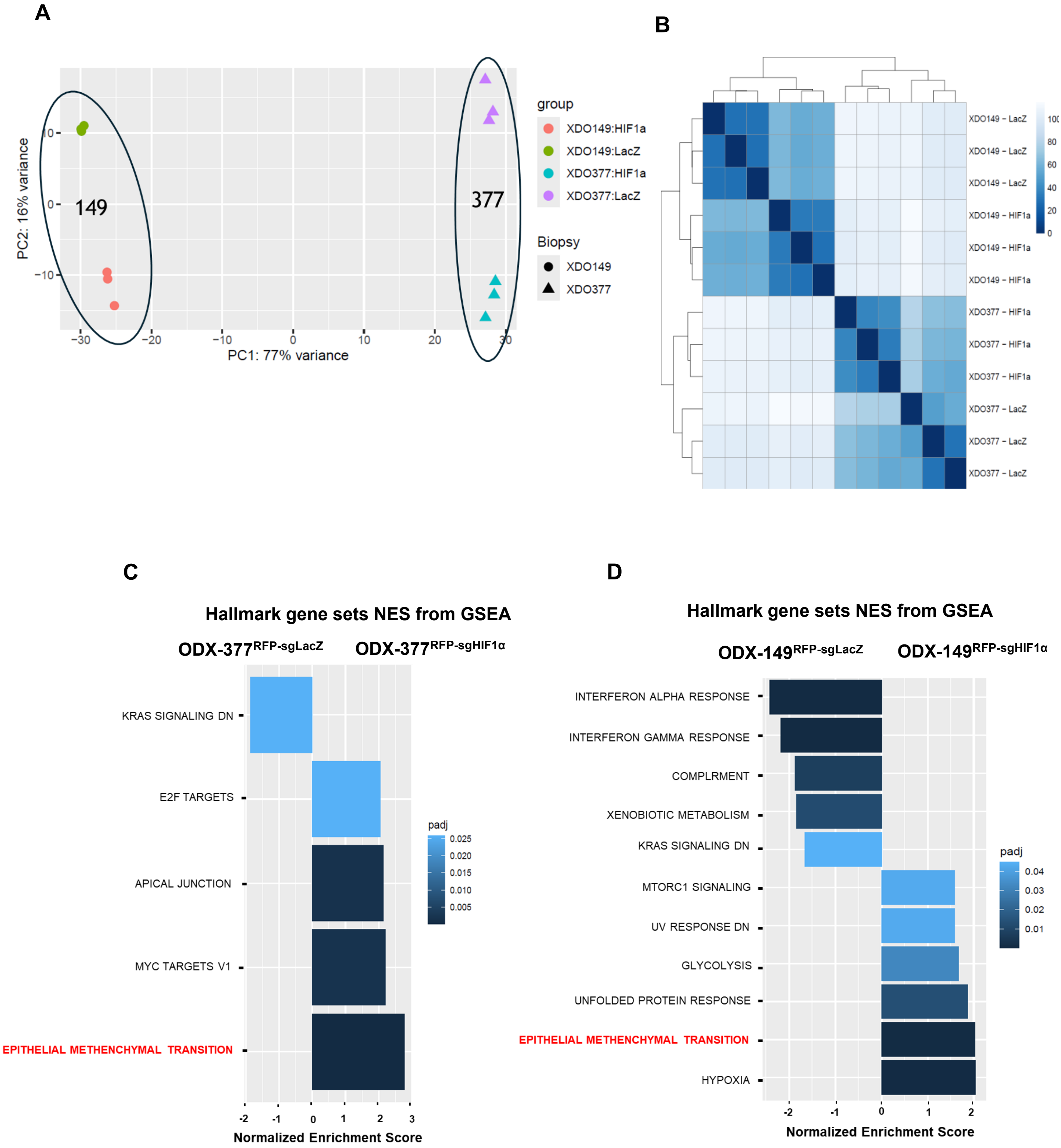


**Supplementary Figure 7. RNA-seq between ODX-377$^{RFP\text{-}sgHIF1\alpha}$/-149$^{RFP\text{-}sgHIF1\alpha}$ and ODX-377$^{RFP\text{-}sgLacZ}$/-149$^{RFP\text{-}sgLacZ}$.** (A) PCA plot among ODX-377$^{RFP\text{-}sgHIF1\alpha}$, ODX-149$^{RFP\text{-}sgHIF1\alpha}$, ODX-377$^{RFP\text{-}sgLacZ}$, and ODX-149$^{RFP\text{-}sgLacZ}$. (B) Sample distance among ODX-377$^{RFP\text{-}sgHIF1\alpha}$, ODX-149$^{RFP\text{-}sgHIF1\alpha}$, ODX-377$^{RFP\text{-}sgLacZ}$, and ODX-149$^{RFP\text{-}sgLacZ}$. (C) Gene set enrichment analysis using Hallmark between ODX-377$^{RFP\text{-}sgLacZ}$ vs ODX-377$^{RFP\text{-}sgHIF1\alpha}$. (D) Gene set enrichment analysis using Hallmark between ODX-149$^{RFP\text{-}sgLacZ}$ vs ODX-149$^{RFP\text{-}sgHIF1\alpha}$.

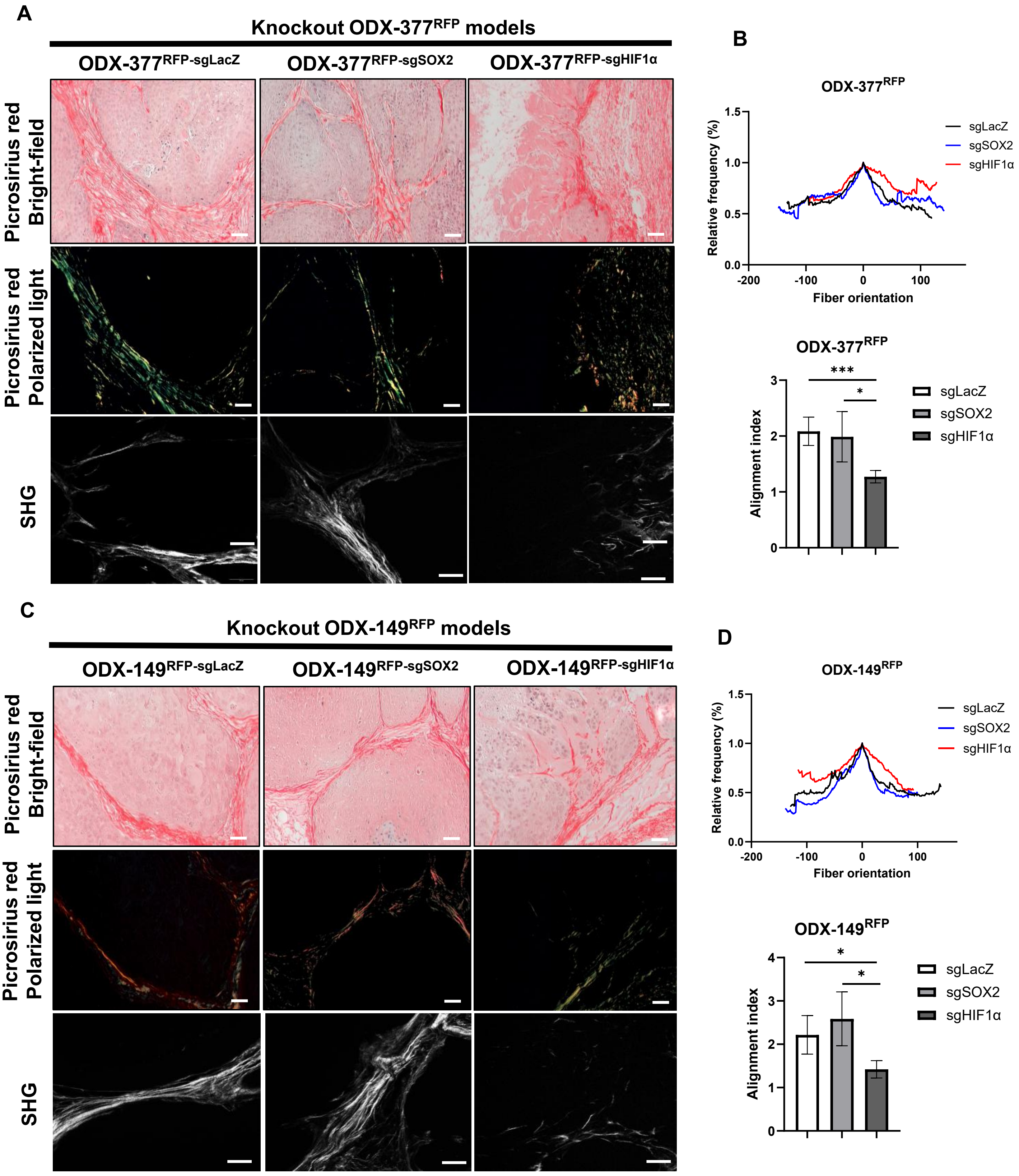


**Supplementary Figure 8.** (A) Picrosirius red staining in ODX-377$^{RFP\text{-}sgLacZ/\text{-}sgSOX2/\text{-}sgHIF1\alpha}$. (B) The fiber orientation and alignment index in ODX-377$^{RFP\text{-}sgLacZ/\text{-}sgSOX2/\text{-}sgHIF1\alpha}$ are shown. Broad distributions indicate low alignment, whereas sharp peaks indicate high alignment. Reduced collagen alignment was observed in ODX-377$^{RFP\text{-}sgHIF1\alpha}$. (C) Picrosirius red staining in ODX-149$^{RFP\text{-}sgLacZ/\text{-}sgSOX2/\text{-}sgHIF1\alpha}$. (D) The fiber orientation and alignment index in ODX-149$^{RFP\text{-}sgLacZ/\text{-}sgSOX2/\text{-}sgHIF1\alpha}$ are shown. Reduced collagen alignment was observed in ODX-149$^{RFP\text{-}sgHIF1\alpha}$. The red color in the bright-field images indicates collagen fibers. The yellow color in the polarized light images indicates collagen type I. The green color in the polarized light images indicates collagen type III. Scale bars are 50 μm (A) (C).

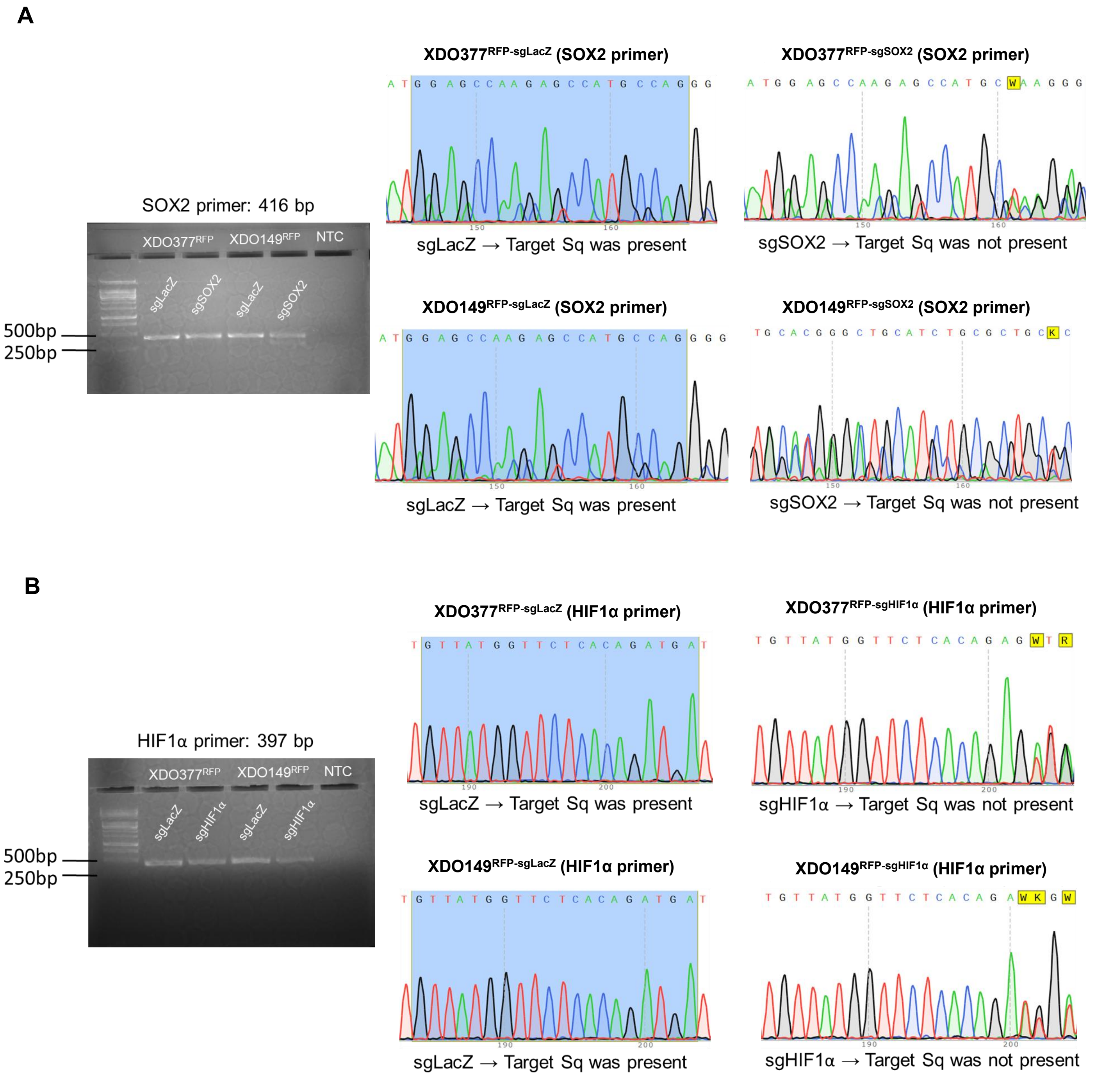


**Supplementary Figure 9. DNA sequencing of genomic DNA to confirm CRISPR-Cas9-mediated knockout.** (A) The PCR product was verified by gel electrophoresis. Sanger sequencing of XDO377$^{RFP\text{-}sgSOX2}$ and XDO149$^{RFP\text{-}sgSOX2}$ is shown, with the blue highlight indicating the target sequence of CRISPR/Cas9-mediated knockout. In the LacZ knockout models, the target sequence is intact, whereas in the SOX2 knockout models, part of the target sequence is deleted. (B) XDO377$^{RFP\text{-}sgHIF1a}$ and XDO149$^{RFP\text{-}sgHIF1a}$ were also confirmed the function of CRISPR-Cas9 using the same method as in (A).

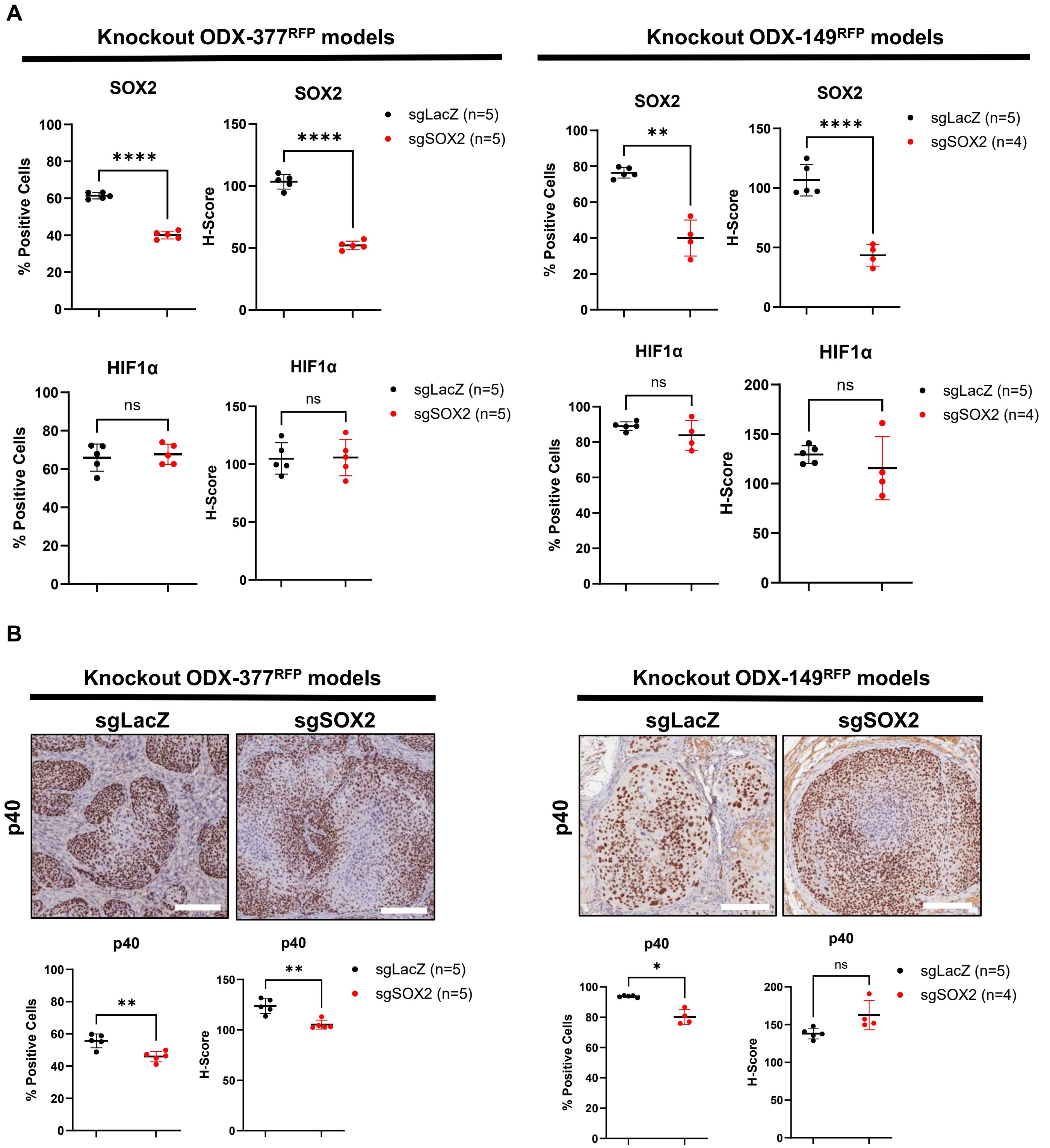


**Supplementary Figure 10.** (A) The staining intensity in Figure 4E was quantitatively evaluated using % positive stained cells and H-score with HALO software. (B) IHC staining for p40 between ODX-377$^{RFP\text{-}sgLacZ}$/-149$^{RFP\text{-}sgLacZ}$ vs ODX-377$^{RFP\text{-}sgSOX2}$/-149$^{RFP\text{-}sgSOX2}$. Scale bars are 200 μm. The staining intensity was quantitatively evaluated using % positive stained cells and H-score with HALO software.

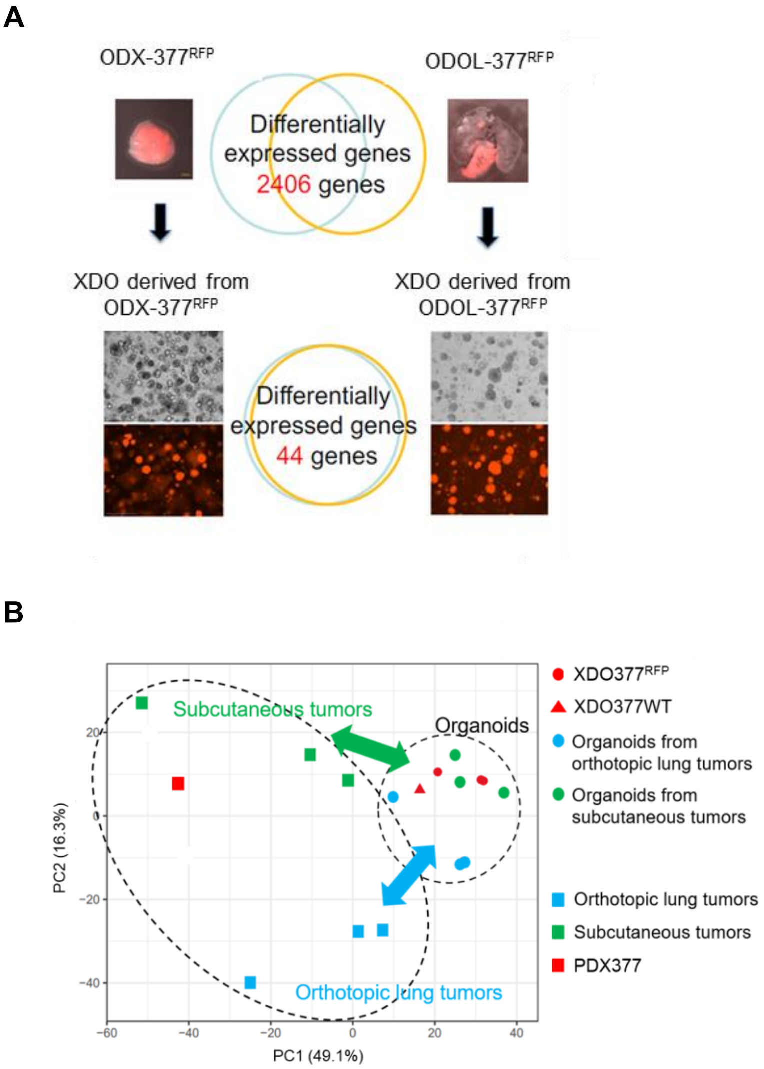


**Supplementary Figure 11.** (A) Organoids were re-established from ODX-377$^{RFP}$ and ODOL-377$^{RFP}$. DEGs were assessed among ODX-377$^{RFP}$, ODOL-377$^{RFP}$, and re-established organoids. (B) The principal component analysis was performed among ODX-377$^{RFP}$, ODOL-377$^{RFP}$, re-established organoids, and original organoid XDO377$^{RFP}$.

**A**

**HPCA**

| HPCA celltype | ODOL | ODX | ORG |
|---|---|---|---|
| Epithelial_cells | 57% | 74% | 92% |
| Keratinocytes | 3% | 8% | 1% |
| MSC | 1% | 1% | 0% |
| Neuroepithelial_cell | 1% | 1% | 0% |
| unclassified | 38% | 16% | 6% |
| All | 100% | 100% | 100% |

**HLCA**

| HLCA celltype | ODOL | ODX | ORG |
|---|---|---|---|
| AT1 | 3% | 8% | 10% |
| AT2 | 1% | 0% | 0% |
| Basal | 17% | 45% | 56% |
| Rare | 1% | 0% | 9% |
| Secretory | 14% | 28% | 10% |
| unclassified | 64% | 19% | 15% |
| All | 100% | 100% | 100% |

**B**

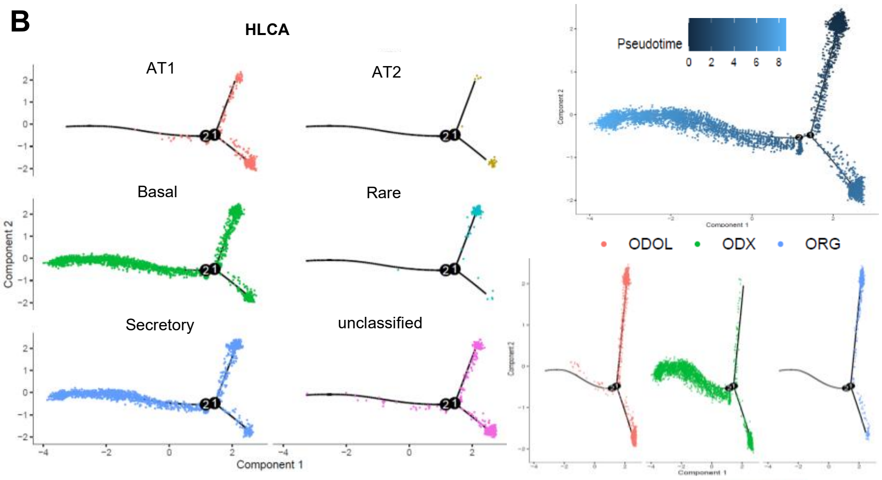


**C**

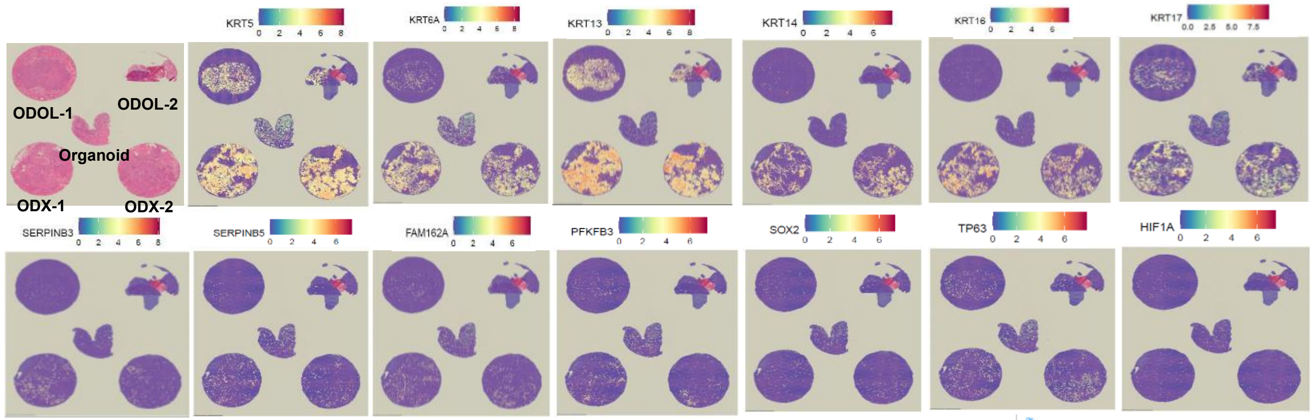


**D**

**HPCA**

| HPCA celltype | ODOL | ODX | ORG |
|---|---|---|---|
| Epithelial_cells | 26% | 20% | 96% |
| Keratinocytes | 11% | 72% | 3% |
| MSC | 3% | 0% | 0% |
| Neuroepithelial_cell | 1% | 0% | 0% |
| unclassified | 59% | 8% | 1% |
| All | 100% | 100% | 100% |

**HLCA**

| HLCA celltype | ODOL | ODX | ORG |
|---|---|---|---|
| AT1 | 23% | 5% | 15% |
| AT2 | 20% | 0% | 0% |
| Basal | 15% | 66% | 78% |
| Rare | 0% | 0% | 2% |
| Secretory | 1% | 10% | 0% |
| unclassified | 40% | 18% | 5% |
| All | 100% | 100% | 100% |

**E**

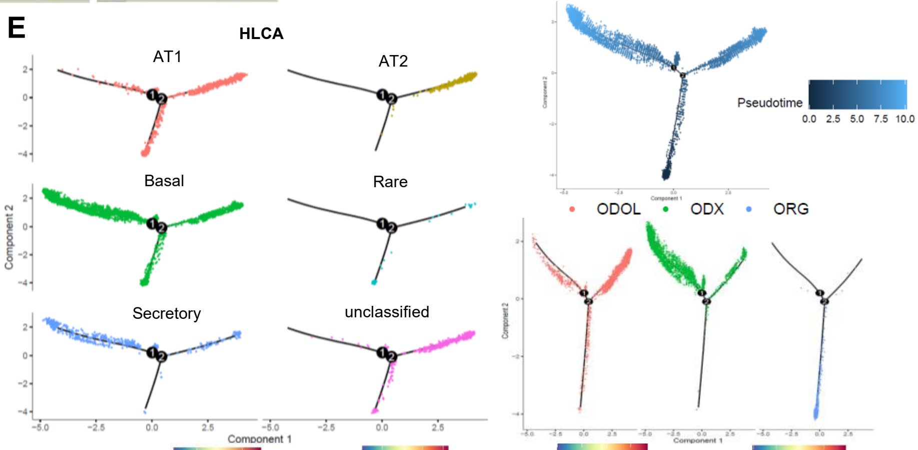


**F**

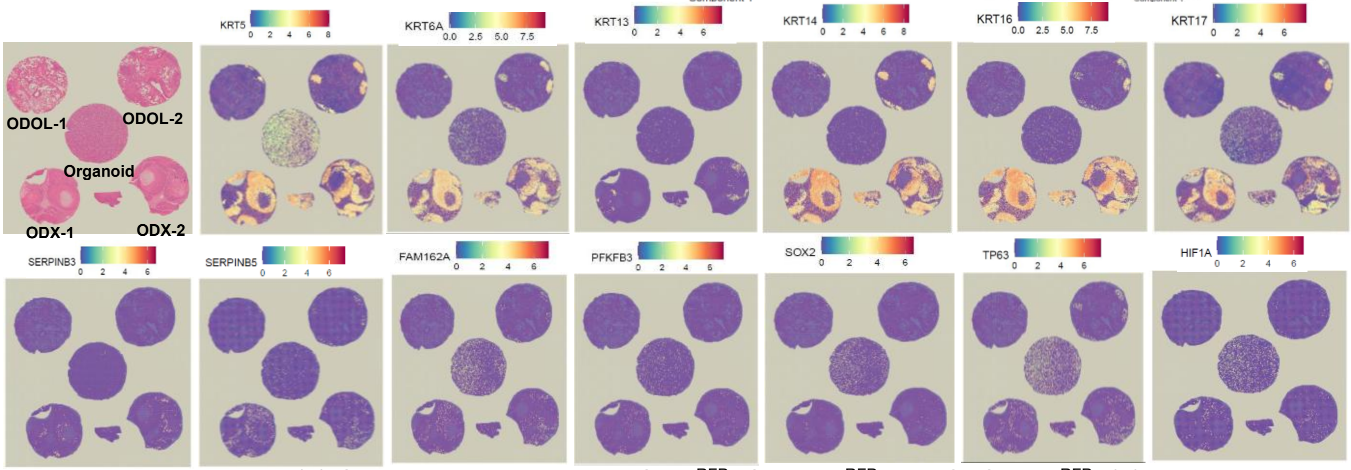


**Supplementary Figure 12.** (A) Cluster analysis among $XDO377^{RFP}$, $ODX\text{-}377^{RFP}$, and $ODOL\text{-}377^{RFP}$. (B) Pseudotime analysis among $XDO377^{RFP}$, $ODX\text{-}377^{RFP}$, and $ODOL\text{-}377^{RFP}$. (C) RNA expression among $XDO377^{RFP}$, $ODX\text{-}377^{RFP}$, and $ODOL\text{-}377^{RFP}$. (D) Cluster analysis among $XDO149^{RFP}$, $ODX\text{-}149^{RFP}$, and $ODOL\text{-}149^{RFP}$. (E) Pseudotime analysis among $XDO149^{RFP}$, $ODX\text{-}149^{RFP}$, and $ODOL\text{-}149^{RFP}$. (F) RNA expression among $XDO149^{RFP}$, $ODX\text{-}149^{RFP}$, and $ODOL\text{-}149^{RFP}$.
